# A Cellular Reference Resource for the Mouse Urinary Bladder

**DOI:** 10.1101/2021.09.20.461121

**Authors:** Dylan Baker, Iman M. Al-Naggar, Santhosh Sivajothi, William F. Flynn, Anahita Amiri, Diane Luo, Cara C. Hardy, George A. Kuchel, Phillip P. Smith, Paul Robson

**Author notes:** These authors contributed equally to this work (Dylan Baker, Iman M. Al-Naggar).

## Abstract

The urinary bladder functions as a reservoir to store and extrude liquid bodily waste. Significant debate exists as to this tissue’s cellular composition and genes associated with their functions. We use a repertoire of cell profiling tools to comprehensively define and spatial resolve cell types. We characterize spatially validated, basal-to-luminal gene expression dynamics within the urothelium, the cellular source of most bladder cancers. We define three distinct populations of fibroblasts that spatially organize from the sub-urothelial layer through to the detrusor muscle, clarifying knowledge around these controversial interstitial cells, and associate increased fibroblasts with aging. We overcome challenges of profiling the detrusor muscle, absence from earlier single cell studies, to report on its transcriptome with many novel and neuronal-like features presumably associated with neuromuscular junctions. Our approach provides a blueprint for tissue atlas construction and the data provides the foundation for future studies of bladder function in health and disease.

## INTRODUCTION

The urinary bladder is a complex organ with diverse cell types interacting in concert to achieve effective storage and expulsion of urine. The cells of the bladder are structured into three major layers; mucosa, submucosa and muscularis. The mucosa is composed of a specialized epithelium - the urothelium - which provides a barrier preventing urine from re-entry into the body, undergoes significant morphological changes to accommodate the dynamics in urine volume, and participates in defense against bacterial infection. The submucosa is an extracellular (ECM) and fibroblast rich layer joining the muscularis and mucosa which has been postulated as an integrative hub of bladder activity. Finally, the muscularis is composed primarily of the detrusor smooth muscle which relaxes and contracts to accommodate and expel urine. The bladder is highly innervated, though lacking neuronal cell bodies itself, and the entire bladder structure considered to be an integrated sensory web that promotes reciprocal communication between the urothelium, the underlying submucosa and muscularis, and the CNS^1^.

Obfuscation of the foundational cellular elements of the bladder has made it difficult to address bladder related disorders leading to common debilitating and costly clinical conditions. Immune and urothelial subtypes are heavily implicated in divergent treatment outcomes and disease recurrence for interstitial cystitis and bladder cancer. Yet the diversity and dynamics of these subtypes is still being investigated with some aspects remaining controversial. Similarly, age-related bladder disorders, especially overactive bladder disorder, have been associated with multiple cell types including the detrusor smooth muscle, fibroblasts and the urothelium. However, contradictory evidence exists with regards to subtype contributions to disease and changes in cell type proportions with aging. Furthermore, a network of bladder interstitial cells, analogous to the interstitial cells of Cajal (ICC) in the gut has been postulated as a critical bladder regulatory element^2–4^. However, definitive characterization of such cells is lacking with conflicting evidence for a unique urinary ICC cell type leading to questions regarding their existence and proposed functionality in the bladder^4^. Thus, it is imperative to provide more comprehensive definitions of bladder cell types in order to better understand pathophysiology and improve patient outcomes.

With the goal of addressing controversies in the field that have arisen due to often incomplete and inappropriate cell typing data, we set out to generate a comprehensive cell atlas of the mouse urinary bladder by combining a set of cell profiling technologies. Here we provide an atlas of the mouse urinary bladder utilizing single cell RNA sequencing (scRNAseq), single nucleus RNA sequencing (snRNAseq), RNA sequencing on bulk tissue, spatial transcriptomics (ST), and imaging mass cytometry (IMC). Using the complementary nature of these techniques we are able to provide novel insight into bladder biology including differentiation of the urothelium, the identity of interstitial cells, and the unique nature of the detrusor smooth muscle.

## RESULTS

### Construction of the Atlas

To provide a comprehensive atlas of the mouse urinary bladder we utilized a number of high content, complementary techniques to generate molecular and spatial profiles for each cell type (Fig. 1A). In-house scRNAseq data was generated from eighteen mouse bladders in five sets utilizing the droplet-based Chromium system in which 3’-end counts were generated. We then combined in-house generated data with publicly available data from the Mouse Cell Atlas^5^ and the Tabula Muris Consortium^6^. To identify potential biases introduced through tissue dissociation and cell capture, we performed differential expression analysis between bulk RNA-seq generated from whole mouse bladders and the aggregate scRNAseq data. Genes found to be significantly over-represented in bulk whole bladder samples were associated with smooth muscle (*e.g. Acta2*, *Actg2*, *Myh11*) and neuronal (*e.g. Gria1*, *Cdh8*, *Rbfox3, Rims1*) cell types (Table S2). A number of these genes (*Hcn1*, *Stum*, *Kcnf1*) were entirely absent from processed scRNAseq data (Table S3). The mouse bladder lacks neuronal cell bodies and thus we postulated these apparent neuronal transcripts were derived from RNAs present within axonal boutons, which would be lost upon tissue dissociation. Of greater concern was the dramatic loss of smooth muscle transcripts. For instance, *Myh11* and *Myocd,* both classical smooth muscle transcripts required for proper smooth muscle function were abundantly expressed in bulk RNA-seq data (*Myh11* 11.6 CPM, *Myocd* 4.42 CPM) with high log fold change (*Myh11* -9.42, Myocd -7.95) in the scRNAseq data^7, 8^. An analysis of the initial scRNAseq data set indicated a population of smooth muscle-like cells but only representing 1.1% of the total, a significant under-representation considering the detrusor muscle comprises a major component of the bladder. The associated cells had high expression of genes known to mark vascular smooth muscle cells (vSMC; e.g. *Pln*) and pericytes (*e.g. Rgs5*), but minimal expression of *Myh11* and *Myocd*. vSMCs and pericytes are readily captured by this scRNAseq approach in other tissues^9, 10^ and are expected to be present in the vascularized bladder. We therefore reasoned that, like cardiomyocytes in the heart^9^, detrusor smooth muscle (DSM) cells do not survive standard dissociation approaches and therefore their transcriptomes missed in our initial, and the previously published^5, 6^ data sets.

**Figure 1.**
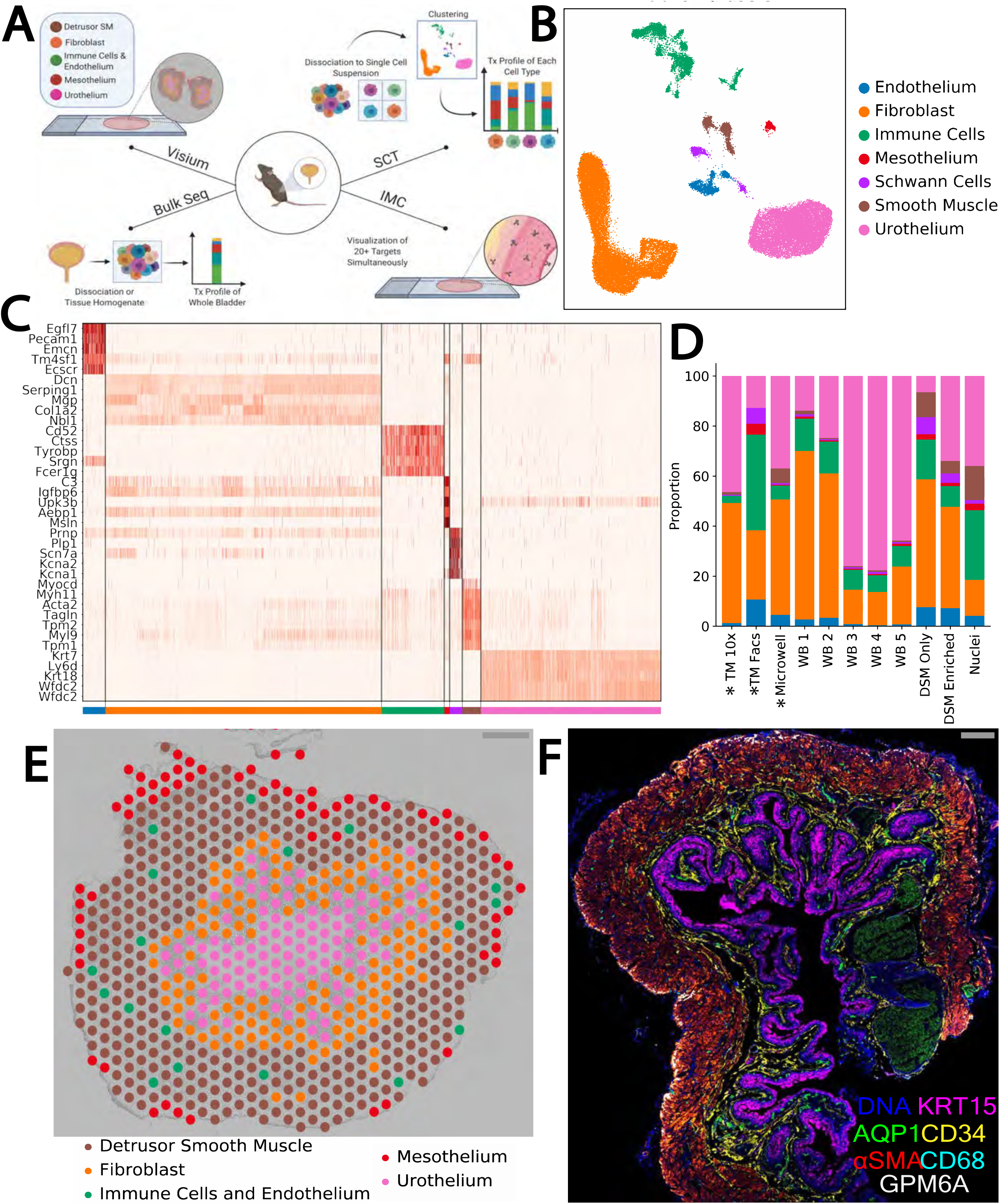
Generation of a Mouse Bladder Atlas via Complementary Dissociative and Spatially Resolved Techniques. A) Schematic of atlas generation design representing the four complementary techniques employed; single cell (and nucleus) transcriptomics (SCT), bulk RNA sequencing (Bulk Seq), imaging mass cytometry (IMC), and spatial transcriptomics (Visium). B) UMAP plot of combined, complete scRNAseq datasets of in house-generated and publicly available datasets totaling 42,904 cells across 11 datasets. C) Gene expression heatmap of select classical and novel marker genes identified for the seven major clusters represented in B. D) Proportional representations of cells within clusters for each individual dataset. Asterisks indicate public datasets. E) Representative image of Visium spatial transcriptomics showing spatial localization of cell types. F) Representative IMC image of mouse urinary bladder. Seven of 15 channels are represented corresponding to the three major layers (mucosa:KRT15, submucosa:CD34, muscularis:αSMA), vasculature (AQP1), myeloid cells (CD68), and the outer mesothelial layer (GPM6A).

To rectify the extreme under-representation of DSM cells we performed additional scRNAseq on samples from optimized dissociation techniques specifically designed to capture DSM cells^11^ by stripping the detrusor layer from the mucosa and processing the detrusor layer with papain followed by collagenase. We also performed snRNAseq which has been previously shown to be useful in capturing difficult to dissociate cell types^9^. The addition of detrusor muscle layer-enriched datasets increased the total smooth muscle cell number from 315 to 1763, and a distinct smooth muscle subcluster became apparent which highly expressed genes originally enriched in the bulk RNAseq over the scRNAseq (including *Myh11* and *Myocd*) and known to be expressed and functional in detrusor muscle (see below), thus providing adequate representation of the detrusor muscle. A complete list of mice, preparation techniques and metrics for each sample in the final dataset can be found in Table S1 and Methods.

With the addition of this detrusor-enriched scRNAseq data, the final single cell transcriptome dataset consisted of a total of 42,904 cells of which 38,858 cells were generated in this study, and represented a gain of 5,296 genes detected compared to the previously published studies. The final dataset correlated well with bulk data (scRNAseq versus whole tissue R=0.915, scRNAseq versus dissociated cells R=0.923; Fig. S1C&D). Primary clustering revealed seven broad clusters corresponding to, in decreasing order of abundance, fibroblast, urothelial, immune, smooth muscle, endothelial, Schwann, and mesothelial cells (Fig. 1B&C). Cell type proportions varied across datasets indicating the impact of tissue preparation techniques on cell types identified and their relative abundance (Fig. 1D). Described in detail throughout the remainder of this study, through iterative clustering and analysis, we define 25 transcriptionally distinct cell populations.

To position these cell types within the spatial context of the bladder we utilized IMC and ST. IMC provides a highly multiplexed antibody-based approach to detect epitopes of interest at sub-cellular (1 μm) resolution. The selection of antibodies for use in IMC was informed by the scRNAseq analysis and designed to encompass the cell type composition of the bladder. The ST technology generates spatially resolved polyadenylated transcriptome gene count data with identical molecular biology to the scRNAseq profiling, at a resolution of 55 um (*i.e.* not single cell). ST was performed on eight mouse bladder sections with a total of 4,569 transcriptome data spots yielding an average of 21,821 UMIs and 4,776 genes per spot. These averages varied significantly between layers correlating with total polyadenylated RNA content identified in the ST permeabilization optimization experiment with the urothelial layer showing the highest density of gene expression (Fig. S1B) averaging 45,884 UMIs and 6,832 genes per spot. Clustering of the ST data resulted in the identification of five clusters, four of which corresponded to a particular layer of the bladder: urothelial, lamina propria, detrusor muscle, and the outer mesothelial lining (Fig. 1E). A panel of 15 metal-conjugated antibodies was used in IMC analysis, a representation of six of these is shown in Fig. 1F. The following analysis of the respective layers and cell types integrates the bulk RNAseq, scRNAseq, snRNAseq, IMC, and Visium ST to define the cellular architecture of the mouse urinary bladder.

### Urothelial Cells

Sub-clustering of the urothelium in the scRNAseq data revealed a continuous cluster polarized by classical urothelial markers, *Krt5* for basal and *Upk2* for luminal (Fig. 2A), suggesting these data capture the dynamics of this differentiation. Pseudotime analysis on the entire urothelial cell population was performed in order to generate insight into the transcriptional dynamics of this process. At pseudotime t=0 *Krt5* expression was highest and *Upk2* expression lowest with the reciprocal observed at t=1. *Krt14*, known to mark a subset of *Krt5*+ basal cells^12^, was detectable in a few cells at the basal end of the pseudotime projection. IMC data for KRT5 and PSCA, markers for basal and luminal populations respectively^13, 14^ validated this projection, though the protein expression was much more restricted to either end points than RNA expression profiles (Fig. 2B). We identified a set of genes (e.g. *Krt20*, *Rgs5*, *Fut9*, *Prss27* and *Sprr2a2*; Fig. 2C) with a rapid increase in expression only at later points in pseudotime and reasoned this set might constitute the end-stage transcriptional differentiation into luminal/umbrella cells. To verify this, we took advantage of the ST data by manually annotating ST spots, guided by the H&E image, as basal or luminal based on the positioning of urothelium relative to the bladder lumen (Fig. 2D). We then performed differential expression analysis between basal and luminal cells for both ST and scRNAseq separately and compared top DE genes. The top 50 genes found with increased expression on lumen-associated ST spots showed significant overlap with the late luminal gene set identified in pseudotime analysis of the scRNAseq data (Fig. 2E, TableS9). This combined strategy therefore validates the pseudotime projection and highlights a robust gene set which delineates the most luminal cells.

**Figure 2.**
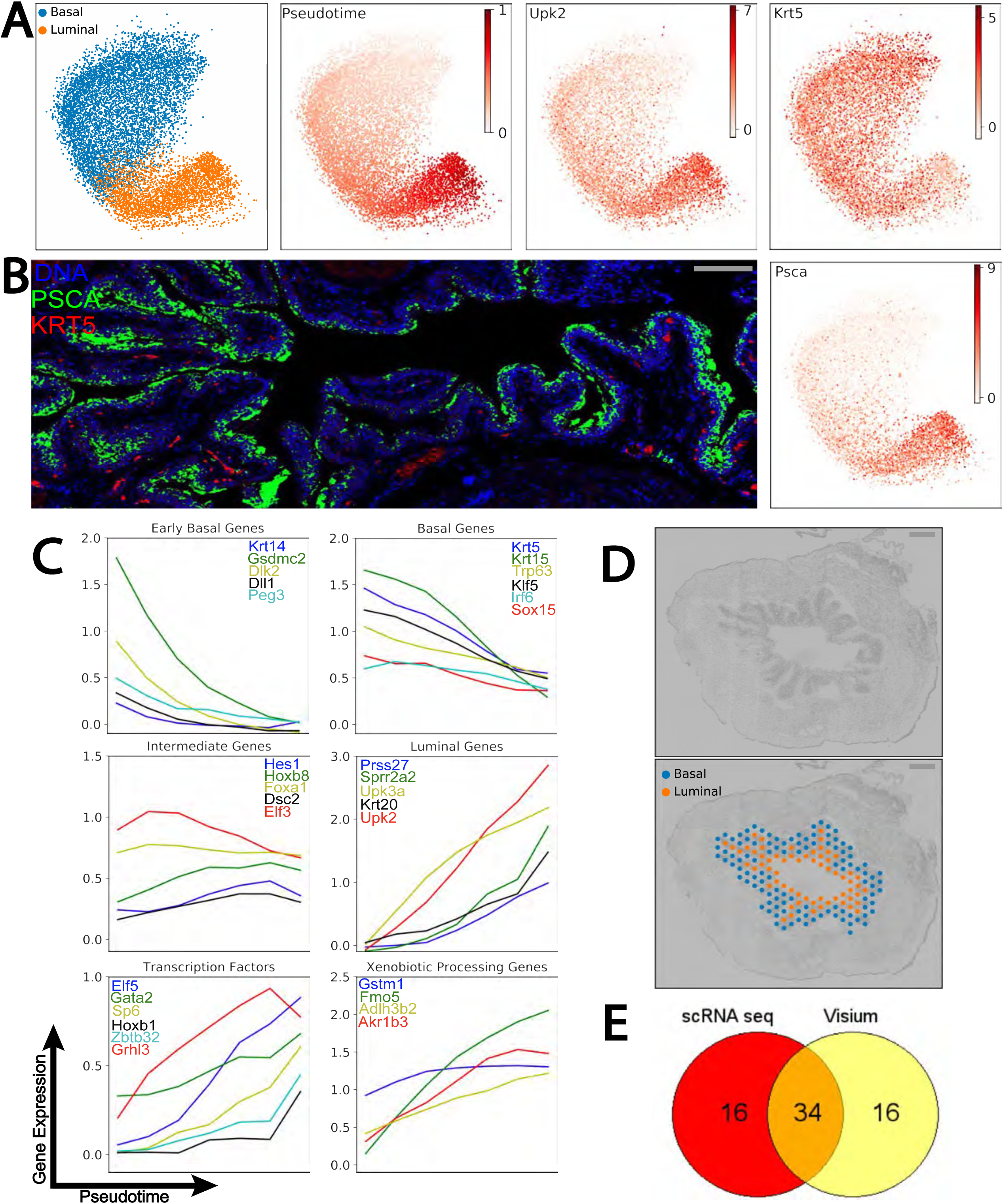
Urothelial Differentiation Dynamics. A) Urothelial cells subclustered and visualized using a Fruchterman Reingold (FR) graph layout reveal two major clusters identified by expression of classical urothelial markers (*Upk2*, *Krt5*). PAGA pseudotime analysis generates a trajectory recapitulating known urothelium differentiation from basal (pseudotime t=0) to luminal (pseudotime t=1) cells. B) IMC (left) of luminal (PscA) and basal (Krt5) markers initially identified by scRNAseq (*Psca* = scRNAseq luminal marker, right) indicates expression discrepancy between transcript (scRNAseq) and protein (IMC) of urothelial markers. C) Pseudotime trajectory plots. Gene expression across discrete points of pseudotime was binned to identify distinct patterns, shown are early basal, basal, intermediate, and luminal. Additional trajectories are shown for selected transcription factors and carcinogen (xenobiotic) processing genes. D) Representative Visium ST image with urothelial spots manually annotated as basal or luminal by spatial location relative to the lumen. E) Venn diagram of top 50 differentially expressed genes identified between basal and luminal cells from manually annotated Visium ST spots (D) and subclustered scRNAseq data.

To further exploit the identified differentiation dynamics we classified sub-groups of genes based on their gradient of expression across urothelial pseudotime (Fig. S2A), defining five general groupings; early basal, basal, intermediate, luminal and late luminal. The pseudotime profiles of genes characteristic for each of these categories is illustrated in (Fig. 2C). Early basal and basal genes decrease across pseudotime with early basal genes showing an exponential decrease in expression and basal genes having a more linear decrease. Similar dynamics are observed between luminal and late luminal for the genes which increase across pseudotime. Intermediate genes had more variable pseudotime profiles, however, each had some increase in expression early in pseudotime with a relative decrease at t=1. The Notch signaling transcriptional effector *Hes1* identified as intermediate in our analysis has previously been shown to transiently increase during urothelial differentiation^15^ lending credence to these genes constituting a true intermediate set. Transcription factors both with known (e.g. *Klf5*^16^) and as yet to be defined (e.g. *SP6*) functions in urothelial differentiation are represented in these gene sets, as are genes (e.g. *Elf3* and *Klf5*) with mutations recently found to be positively selected for in apparently normal human bladders^17^. Interestingly, we identified a group of luminal genes which are each involved in xenobiotic processing based on GO annotations (Fig. 2C). One of these, *Gstm1* is one of the few known loci associated with increased genetic susceptibility to bladder cancer in humans^18^ thus providing a mechanistic link between carcinogens in the bladder lumen and urothelial carcinomas.

### Fibroblasts

Fibroblasts within the lamina propria and the detrusor muscle layer of the bladder wall have been postulated to behave as an intercellular communications network^19–22^ analogous to the interstitial cells of Cajal (ICC) in the gut. However, a uniquely identifiable ICC in the bladder has not been convincingly characterized^3, 23, 24^. Although *Kit* (CD117) is the prototypical marker for ICCs in the intestinal tract^25^, recent data suggested mouse bladder *Kit* is expressed only on mast cells^26^. In our data, *Kit* expression was restricted to a small number of immune and DSM cells and other gut ICC markers (*Pdgfra*, *Entpd2*, *Cd34*, and *Gja1*) were either absent or not restricted to the fibroblast cluster (Fig. 3A).

**Figure 3.**
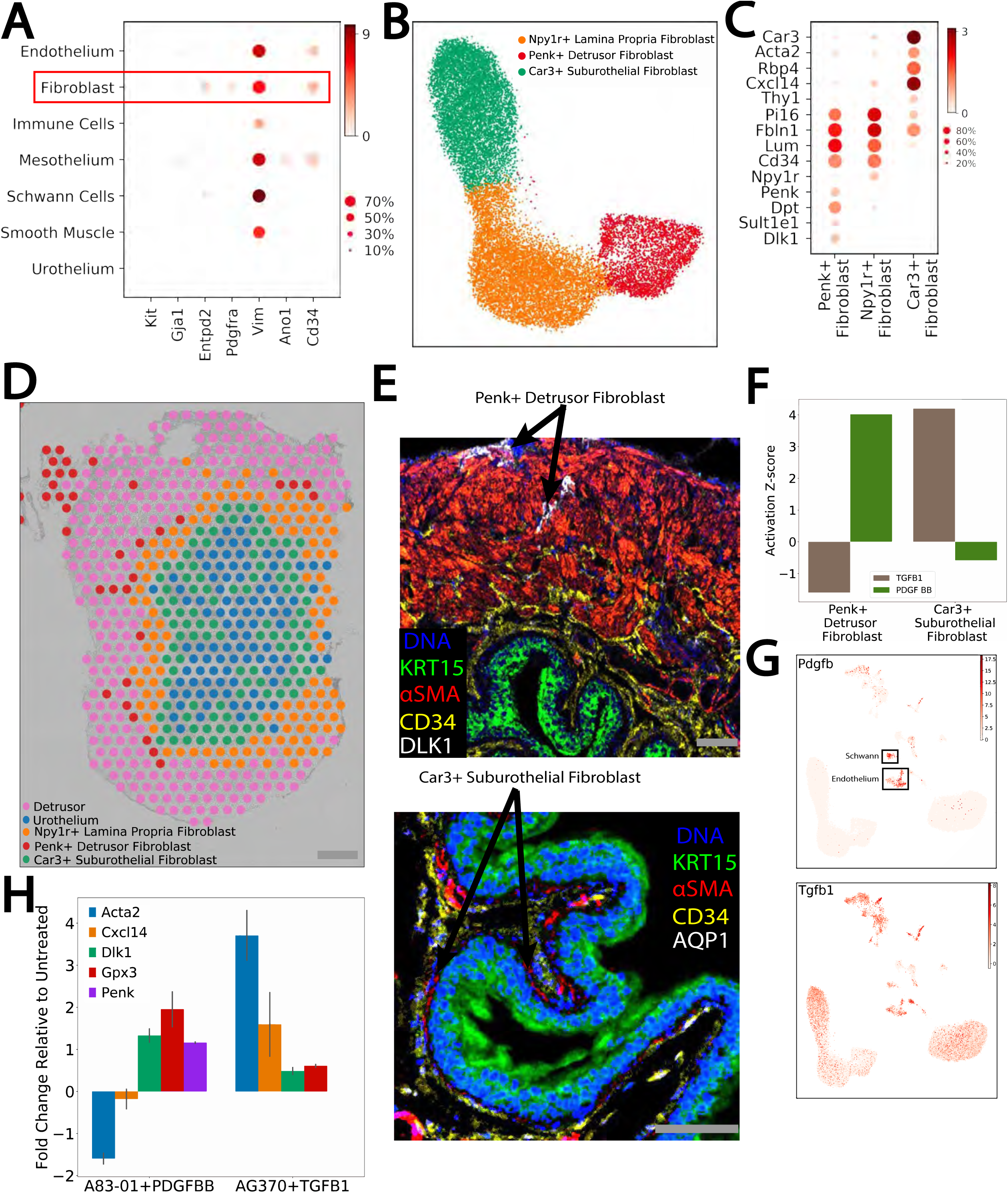
Definition and Spatial Localization of Fibroblast Sub-types. A) Dot plot representing the expression of previously proposed markers of bladder ICC across the seven major cell types in the bladder. B) UMAP plot of fibroblasts defined into three subclusters by the expression of *Car3*, *Npy1r*, and *Penk*. C) Dot plot of markers identified as distinguishing the three fibroblast subtypes identified in B. D) Clustering of ST data based on fibroblast sub-cluster marker genes identified in the scRNAseq data. E) IMC validation of fibroblast localization with Acta2/aSMA+ fibroblasts directly beneath the urothelium (marked by Krt15), and Cd34+ fibroblasts residing primarily in the lamina propria and extending into the detrusor region and Dlk1+ fibroblasts residing in the detrusor region. F) IPA upstream regulator analysis of differentially expressed genes between dmF and suF indicating reciprocal upstream activity (Activation Z-Score) of Tgfb1 and Pdgfbb. H) Quantitative RT-PCR analysis of fibroblast subcluster marker genes, normalized to untreated, from growth factor (Pdgfbb & Tgfb1) and their respective receptor inhibitors (AG370 & A83-01) treated lpF cultures. Primary lpFs were derived derived through FACS and treated for 8 days (suF markers: *Acta2* and *Cxcl14*; dmF markers: *Dlk1*, *Penk*, and *Gpx3*). G) UMAP plot of expression of *Tgfb1* and *Pdgfb* in scRNAseq data.

Further analysis of fibroblasts revealed three sub-clusters (Fig. 3B) each defined by marker genes such as *Car3*, *Npy1r* and *Penk* (Fig. 3C). Utilizing the spatial information provided by ST and IMC, we were able to position these three sub-populations within the bladder layers. The *Penk*+ fibroblasts are primarily located in the detrusor layer (Fig. 3D&E) while the *Npy1r*+ fibroblasts are primarily within the lamina propria creating a network which extends into the detrusor layer (Fig. 3D&E). The *Car3*+ fibroblasts which express myofibroblast-related genes (e.g. *Acta2*) are located directly beneath the urothelium, and are also enriched in basement membrane collagens (*Col4a1*, *Col4a2*). Based on their spatial geography we term these sub-populations as *Car3*+ sub-urothelial (suF), *Npy1r*+ lamina propria (lpF), and *Penk*+ detrusor muscle (dmF) fibroblasts. All appeared to form a continuum with suF co-localizing with lpF in the sub-urothelial region and the terminal ends of the lpF extending into the detrusor layer co-localizing with the dmF (Fig. 3D&E).

Given the spatial and transcriptional proximity between all three fibroblast subtypes we reasoned that the entire population represents different cellular states of a single cell lineage, and that microenvironmental cues may guide transcriptome differences. IPA upstream regulator analysis based on differentially expressed genes between dmF and the suF revealed reciprocal apparent activity of TGFB1 and PDGFBB in these two cell types (Fig. 3F). In scRNAseq data *Tgfb1* was expressed across multiple cell types including the urothelium and the suF (Fig. 3G). *Pdgfb* was expressed highly on a subtype of Schwann cells which primarily reside in the detrusor region (Fig. 3G, 6D&E). ST data showed a higher expression of *Tgfb1* in the urothelial/ lamina propria region compared with the detrusor region while the inverse was true of *Pdgfb* expression (Fig. S2). Thus, the apparent activity of the TGFB1 and PDGFBB pathways in suF and dmF, respectively, fit with the spatial location of the expression of these ligands.

To determine if reciprocal activity of TGFB1 and PDGFBB causes a transition of lpF into suF and dmF, we isolated lpFs by stripping mucosa from detrusor to remove dmFs and collecting the CD34+ mucosal population by FACS. We then cultured the recovered cell population in combinations of these ligands and inhibitors of TGFB1 and PDGFBB pathways. After 6 days in culture the expression of dmF-associated genes (*Dlk1*, *Penk*, *Gpx3*) were increased in PDGFBB activated, TGFB1 inhibited cultures compared to untreated cells (Fig. 3H). Expression of suF-associated genes (*Acta2*, *Cxcl14*) was also decreased in these cultures. In TGFB1 activated, PDGFBB inhibited cultures expression of suF-associated genes were increased compared to untreated samples although dmF-associated genes were also slightly increased. This would indicate that while reciprocal activity of TGFB1 and PDGFBB can influence the transcriptional profile of lpF, additional factors may be involved in the differentiation of the suF population. Notably, these fibroblast sub-populations fit into classifications of cancer-associated fibroblasts that we and others have described in human solid tumors, with the suF favorably comparing with myCAF/CAF-B and dmF to iCAF/CAF-A^27, 28^. ACTA2/SMA-high CAFs have further been identified as a tumor-adjacent TGFβ-driven population^28^, thus correlating with the signaling of the suF positioned adjacent to the urothelium.

### Fibroblasts and Aging

To elucidate the cell types which are responsible for gross changes observed in the aging bladder^29–31^ we analyzed previous bulk data within the context of our scRNAseq data (Fig. 4A). A number of genes found to be upregulated with aging based on previous literature were shown to be expressed in the fibroblast subtypes in scRNAseq data (Fig. 4A). Comparison of cell type proportions from our own data between batch-matched old/young and old/mature datasets showed an 11% increase in the proportion of fibroblasts with age regardless of dissociation technique used (Fig. 4B). Additionally, the ratio of mesenchymal to urothelial cells was found to be increased with aging by flow cytometry (Fig. 4F) indicating an increase in fibroblast number with aging.

**Figure 4.**
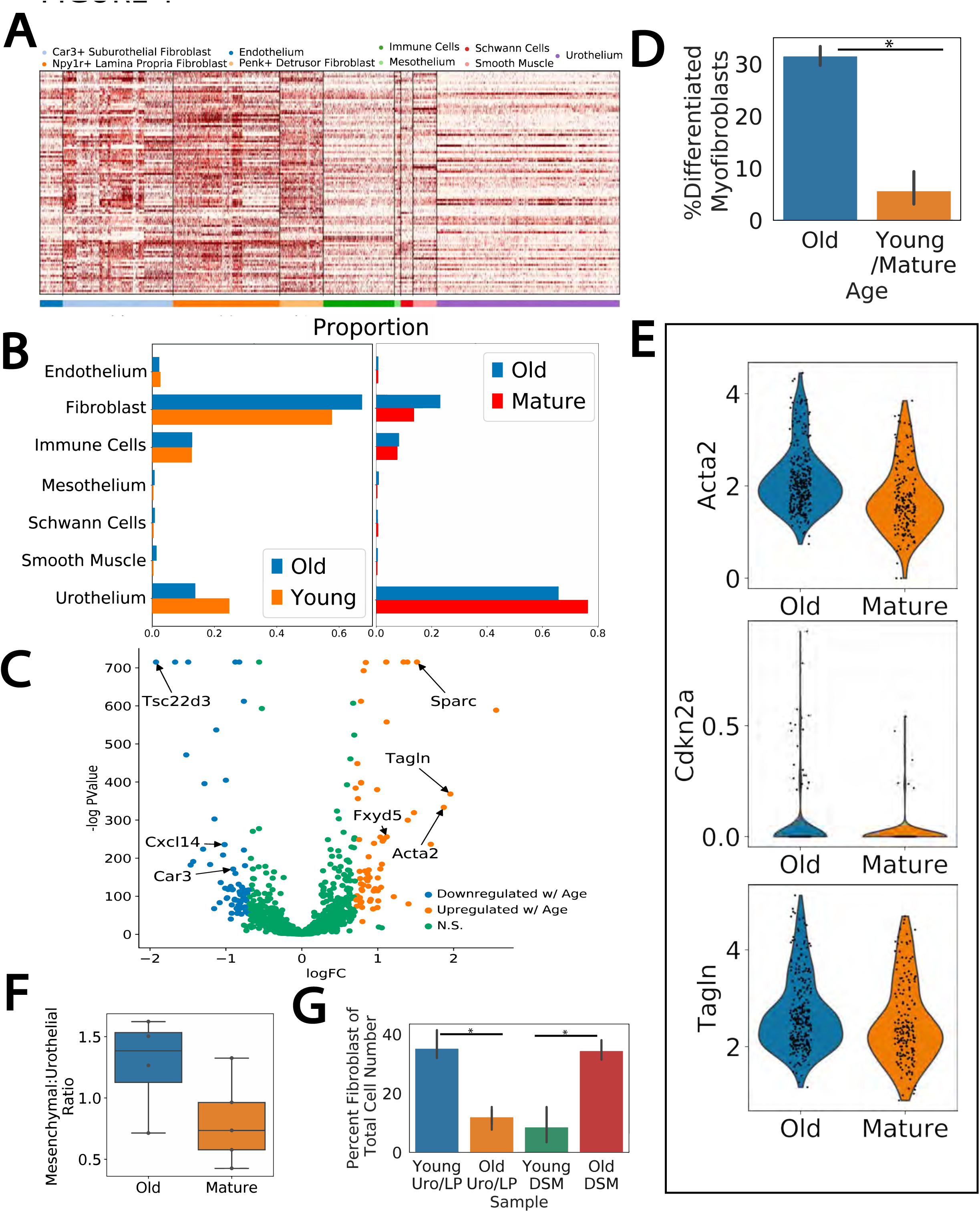
Age Related Changes in Fibroblast Populations. A) A gene expression heatmap across the bladder single cell data set of previously identified genes found to be upregulated in aging studies. Cell types are indicated by color bar. B) Proportion of cell types in scRNAseq data from dissociation matched datasets (WB1+WB2, WB3+WB4+WB5). C) Volcano plot of differential expression analysis between old and young/mature fibroblasts. suF marker genes (*Car3*, *Cxcl14*, *Tsc22d3*) and those related to myofibroblast differentiation (*Sparc*, *Tagln*, *Fxyd5*, *Acta2*) are indicated. D) Percent of suFs classified as terminally differentiated myofibroblasts. E) Violin plot of ST expression of myofibroblast differentiation related genes in suF spots in aged and mature sections. F) Flow cytometry analysis of mesenchymal (VCAM-1+) and urothelial (EPCAM+) populations in old (22M) and mature (12M) bladders. G) Flow cytometry analysis of CD34+ fibroblast populations in the urothelium/lamina propria (Uro/LP) versus the detrusor layer (DSM) in young (4M) and old (22M) bladders. Percent of fibroblasts in each layer significantly changed with aging (Uro/LP p=0.004, DSM p=0.003).

Our findings were in contrast to a recent study utilizing scRNAseq, that found the mesenchymal compartment decreased by a factor of three with aging and the urothelial compartment increased by a similar amount.^32^ As this consortium performed scRNAseq on stripped bladder mucosa (non-mucosa was discarded) rather than whole bladder we reasoned the aged fibroblast, and its associated deposition of ECM, may influence the physical properties of the tissue resulting in differential cell composition in mechanically stripped bladders compared to whole bladder preps. To test this hypothesis, we performed flow cytometry on young and old stripped bladders to determine how the number of fibroblasts change with aging in the two regions, the mucosa (urothelium and lamina propria) and the detrusor/muscularis, after stripping. In young the proportion of fibroblasts was higher in the mucosa than in the detrusor region while the inverse was true of the old samples (Fig. 4F). Thus, the change in physical properties of the bladder with aging alters the composition of the stripped mucosal layer, with more fibroblasts remaining on the surface of the muscularis layer after stripping of the aged bladder than that of the young.

As we have defined three sub-types of fibroblasts we next sought to determine if aged-related changes were sub-type specific. The increases in fibroblast numbers were similar across the three fibroblast subtypes (Fig. 4B). However, differential expression between old and young/mature fibroblasts revealed a number of differentially expressed genes were specific to the suF (Fig. 4C). Many of the suF-specific genes found upregulated in aging had been previously implicated in myofibroblast differentiation (*Tagln*, *Acta2, Sparc*, *Fxyd5*), suggesting the increase in the suF cell number with aging coincides with an increase of suF differentiation towards terminal myofibroblasts. To confirm this we used a set of terminally differentiated myofibroblast markers (*Acta2*, *Tagln, Mylk, Myl9*) to binarize the suF cluster into partially differentiated myofibroblasts and terminally differentiated myofibroblasts by Gaussian mixture modeling, and compared the results across ages. Indeed, the percentage of terminally differentiated suF myofibroblasts increased from ∼6% in both young and mature samples to∼32% in the aged samples (Fig. 4D, p indistinguishable from 0). This was further confirmed by spatial transcriptomics: ST spots annotated as suF had increased expression of myofibroblast differentiation genes *Acta2*, *Tagln*, and *Cdkn2a* when comparing old to mature tissue sections (Fig. 4E). Additional genes emerging in the differentiated myofibroblast, identified by differential expression between the terminally and partially differentiated myofibroblast cells, revealed a number of genes related to neurogenic bladder pathology (Table S5). Specifically, genes such as *Ngf*, *Bdnf*, and *Nrg1* were found to be increased in terminal myofibroblasts and constitute molecular markers for neurogenic bladder syndrome and outlet obstruction^33, 34^.

### Detrusor Smooth Muscle

After ensuring optimization of tissue harvest for DSM-specific cells, the smooth muscle population in the bladder could be sub-divided into three distinct clusters (Fig. 5A) consisting of the 555 vSMCs, 388 pericytes, and 820 DSM (Fig. 5A). Notably, of the cells from the previously published data sets, only 4 cells clustered with the DSM, emphasizing the importance of our optimized tissue enrichment and dissociation of this cell type. Overall, each of the three subclusters shared expression of many general smooth muscle marker genes (e.g. *Acta2*, *Myl9, Mylk*) (Fig. 5B, TableS6). The two vascular-associated populations shared some known molecular markers, such as *Epas1, Sncg*, and *Notch3*^35^, that were absent in the DSM, but also each had distinct molecular features that enabled their classification separately as pericytes (*Rgs5, Kcnj8*, *Pdgfrb*)^36^ and vSMC (*Tesc*, *Pln, Wtip*)^37^. In addition to the earlier-mentioned *Myh11* and *Myocd* expression, the DSM cluster was marked by the mRNA expression of a number of structural proteins associated with the detrusor muscle such as *Actg2*, *Acta1*, and *Tnnt2*. *Actg2* (υSMA) localization to the detrusor layer was confirmed by IMC (Fig. 5C) while the more broadly expressed *Acta2* (αSMA) was found both in the detrusor layer and around vascular structures (i.e. in vSMC) which colocalize with the endothelial marker AQP1 (Fig. 5C). In addition, a number of genes (*Cnn1*, *Synpo2*, *Actg2*, *Mylk*) genetically linked to detrusor muscle defects in megacystis microcolon intestinal hypoperistalsis syndrome^38, 39^ are also marker genes of DSM cells identified in scRNAseq, further validating the identity of this cluster.

**Figure 5.**
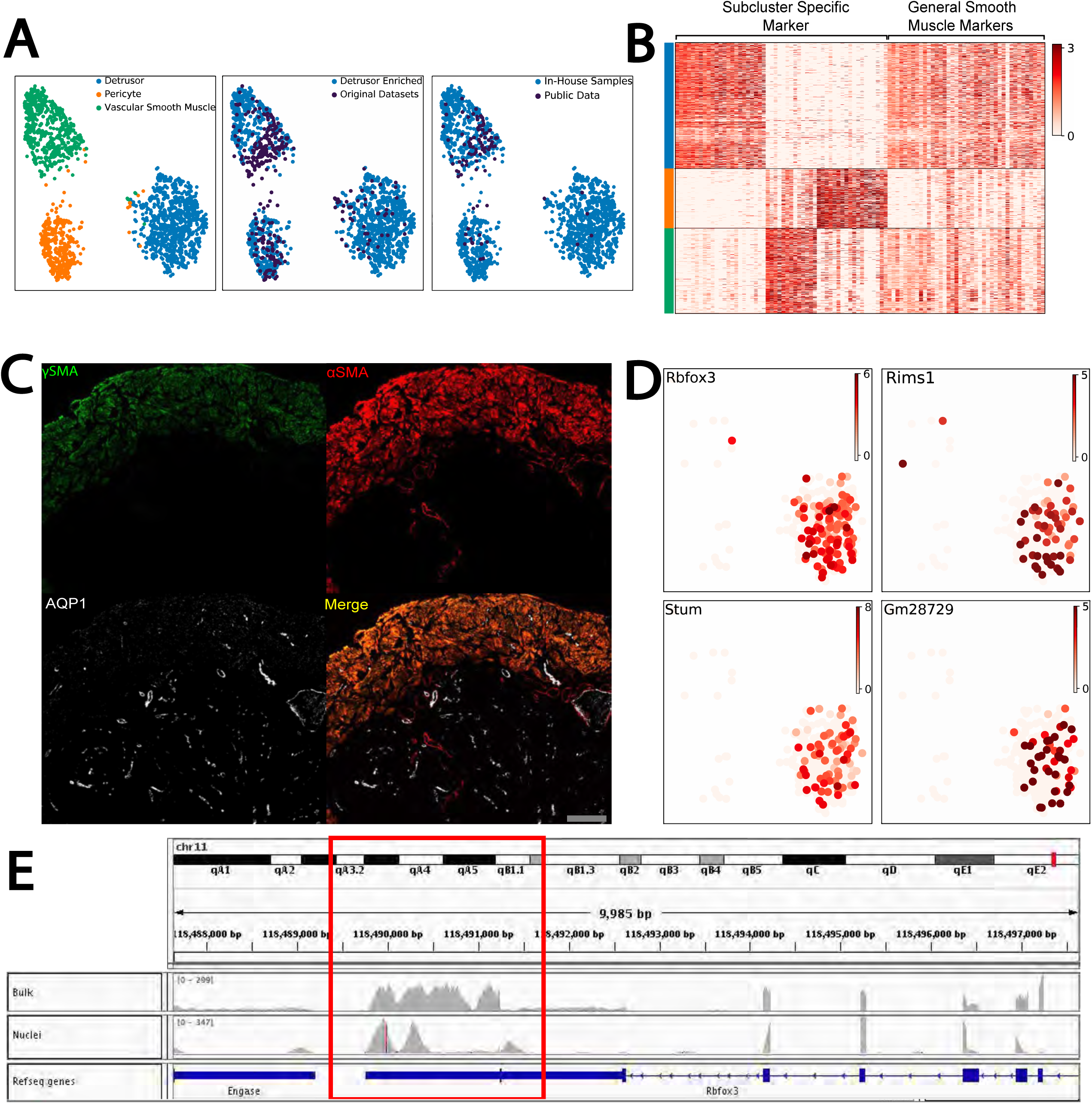
Smooth Muscle Subclustering Identifies a Detrusor Muscle Population. A) UMAP plots of subclustered smooth muscle identified in combined complete dataset color coded by cell type and dataset. B) Gene expression heatmap of smooth muscle marker genes. Subcluster specific genes identified by marker analysis of the subclustered dataset (left) were plotted against the general smooth muscle marker genes identified in the original combined complete dataset. C) IMC validation images of the detrusor smooth muscle subtype identity. υSMA (*Actg2*) was identified as a detrusor cluster specific marker in sequencing data and used as a detrusor marker in IMC. αSMA (*Acta2*), a general smooth muscle marker, localized to the detrusor layer and also co-localized with the endothelial marker Aqp1. D) UMAP plot of smooth muscle subclusters from the snRNAseq dataset. DSM nuclei displayed expression of genes previously indicated as exclusively or primarily neuronal. E) IGV plot of snRNAseq and bulk RNAseq indicate specificity of read mapping to exons of *Rbfox3*.

As previously stated, differential expression between whole bladder bulk RNA-seq and the initial scRNAseq data resulted in a number of genes enriched in the bulk that are classically associated with neuronal cell types (*e.g. Rbfox3*, *Kcnf1, Rims1 (Table S2)).* Initially, we suspected that transcripts of these genes are present within axons and nerve terminals innervating the bladder and thus were lost following dissociation. Interestingly, however, we found the expression of these neuronal genes to be located specifically in the DSM cell cluster. For instance, *Rbfox3*, which encodes NeuN, thought to be exclusively expressed in neurons and is frequently used to mark neuronal nuclei^40^, has not previously been shown to be expressed in smooth muscle cells. With the tight coupling between axonal projections and DSM in the bladder^41^ we initially postulated that the apparent presence of these transcripts in DSM may be due to axonal terminals adhering to DSM cells upon dissociation in the detrusor-enriched preparations. To address this, we analyzed the snRNAseq muscle subclusters by themselves (i.e. without any scRNAseq data) reasoning that any DSM nuclei will only contain transcripts expressed in the DSM cells themselves, and the absence of neuronal nuclei within the bladder itself will remove neuronal-specific transcripts. These neuronal-like transcripts were still however detected and specific to the DSM cluster within the single nuclei data (Fig. 5D). Visualizing these mapped reads confirmed appropriate mapping to the exonic regions of these genes (Fig. 5E) ruling out spurious mapping issues. These data indicate that the DSM has a transcriptional profile related to both the contractile function of detrusor muscles, as well as a transcriptional signature better known in neuronal biology.

To further identify DSM genes that may contribute to bladder-specific biology we refined the *DSM_nocontractile* gene set by applying a bladder specificity index score for each gene utilizing mouse ENCODE transcriptome data (Fig. 6A). A higher index score indicates higher specificity for expression in the DSM compared to all other tissues profiled in the ENCODE data and therefore potentially connotate DSM-specific functionality. The two genes with the highest specificity both encode neuropeptide Y receptors, *Npy6r* and *Npy4r*. As far as we are aware, this is the first report of these receptors on DSM cells, this is intriguing as Npy4r has been identified as expressed on colonic muscle cells and functionally involved in colonic contraction.^42^At least one of the ligands for these receptors, NPY, is known to be richly distributed in nerve fibres within the detrusor layer^43^. *Stum*, first characterized as a mechanosensing molecule in *Drosophila*^44^, is also specific to the DSM among bladder resident cells. An as yet to be functionally characterized gene, *Gm28729*, is also represented in this gene set. Comparative analysis of the *Gm28729* predicted protein sequence indicates a deep evolutionary history (invertebrates through to mammals) of this protein-coding gene, with an apparent functional loss within the last common ancestor of humans and Denisovans (Fig. S3). Thus, numerous genes are revealed in our uncovering of the DSM transcriptome with apparent functional relevance to bladder control potentially through roles at the neuromuscular junction.

**Figure 6.**
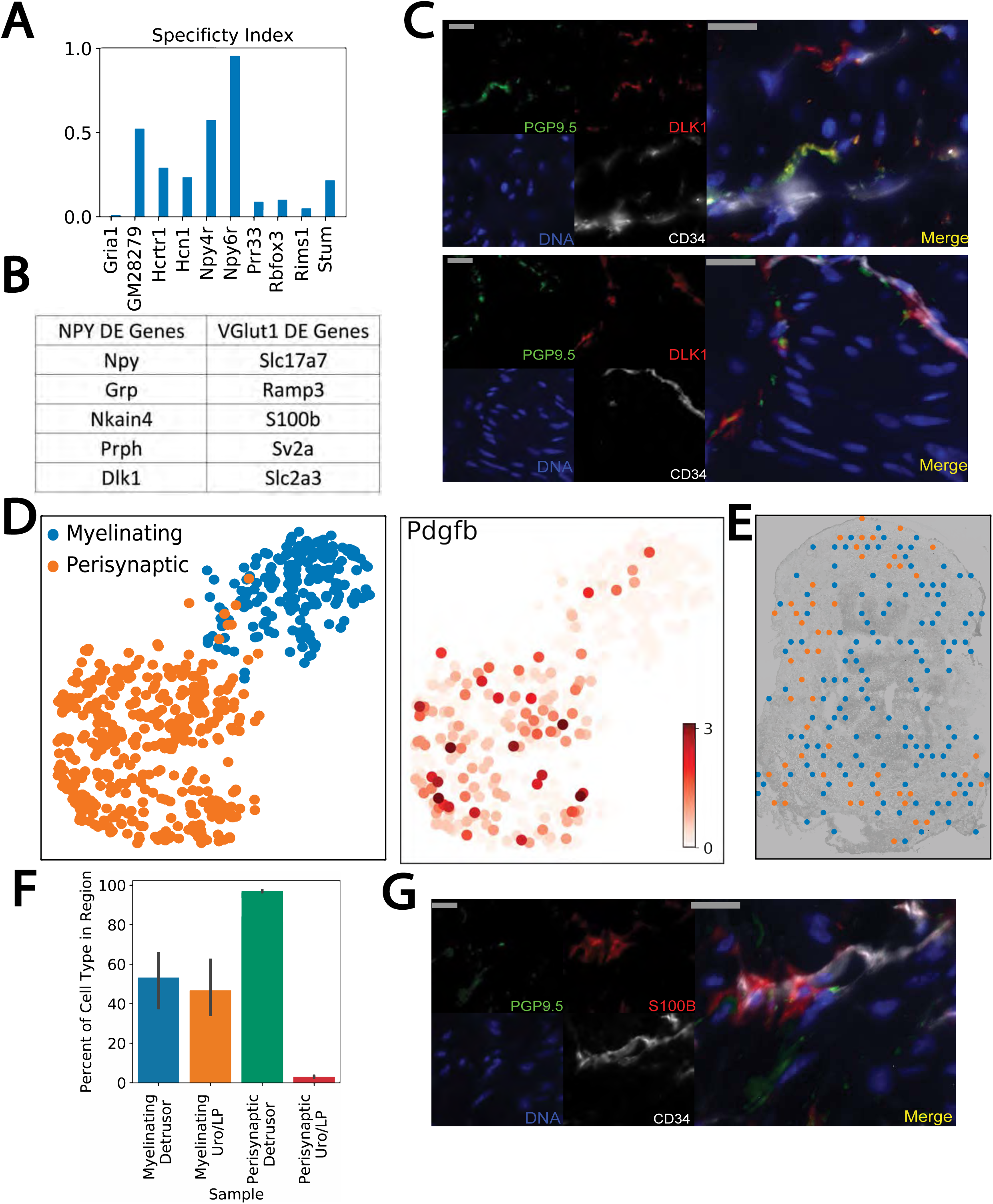
Neuronal Related Features of the Mouse Bladder. A) Bladder specificity of genes differentially expressed in bulk data compared to scRNAseq. Index derived from mouse ENCODE transcriptome data and defined as bladder RPKM over sum RPKM of all ENCODE tissues. B) Differentially expressed genes between *Npy*+ and *Npy*- ST spots (left) and *Slc17a7*+ and *Slc17a7*- ST spots. C) Representative immunofluorescence images of mouse bladder sections co-stained with Pgp9.5, Dlk1, and Cd34. D) Left panel - UMAP plot of subclustered Schwann cells indicating two subclusters, myelinating and perisynaptic. Right panel – the same UMAP plot indicating *Pdgfb* expression. E) Representative ST image of Schwann cell containing spots color coded by perisynaptic or myelinating gene signatures. F) Quantification of Schwann cell subtype localization across all ST sections. G) Representative immunofluorescence images of mouse bladder detrusor region co-stained with Pgp9.5, S100b and Cd34 to highlight close proximity of fibroblasts and Schwann cells within the detrusor region.

### Neuronal Moieties

While no neuronal cell bodies are present in the mouse bladder, we sought to identify transcripts from innervating neurons in ST using genes encoding for products known to localize to neuronal boutons in the bladder; specifically, *Npy* and *Slc17a7* (VGLUT1)^45^. Eight spots were found to contain *Npy* and differential expression between *Npy*+ and *Npy-* spots revealed Npy was co-expressed with other neuronal-associated genes such as *Grp* (Fig. 6B, S4A, TableS7) which has been used as a marker for neuronal processes^46^. Based on this and the fact these genes are absent in our scRNAseq data, including the DSM-enriched samples, provides confidence they represent neuronal-specific transcripts. However, co-expression of *Npy* and *Dlk1* was surprising as *Dlk1* was found to be expressed in the *Penk*+ dmF; co-expression in ST could be due to co-localization of axons with *Dlk1*+ fibroblasts or due to *Dlk1* transcripts within the axons themselves. Previous work has indicated that *Dlk1* is expressed on neurons and in fact distinguishes fast twitch from slow twitch motor neurons^47^. Given the physiological implications of axons with a fast twitch transcriptional signature innervating the bladder we sought to resolve where Dlk1 protein is present. Bladder sections were co-stained with Dlk1, Pgp9.5 and Cd34 to distinguish Dlk1 signal in neurons vs fibroblasts. Dlk1 signal co-localized independently with both Pgp9.5 and Cd34 indicating its expression in both neurons innervating the bladder and *Penk*+ dmFs (Fig. 6C).

Only three *Slc17a7*-containing spots were identified in ST. Differential expression analysis, despite being underpowered and lacking significance, between *Slc17a7*+ and *Slc17a7*- spots revealed *Slc17a7* co-expression with *Ramp3*, *S100b*, *Sv2a* and *Slc2a3*. With the exception of *S100b* each encode for neuronal specific products (Fig. 6B, S4A). *S100b* is a well-known Schwann cell marker and based on the association between Schwann cells and axons this co- expression of *S100b* and *Slc17a7* in ST is intuitive. Sub-clustering of Schwann cells in the scRNAseq data (*S100b*+) resulted in two clusters identified as myelinating and non-myelinating Schwann cells based on classical markers (Fig. 6D). ST indicated that peri-synaptic Schwann cells reside mostly within the detrusor region (Fig. 6E and 6F). Interestingly, scRNAseq data indicated that Schwann cells are a primary source of *Pdgfb* which we determined to be an upstream regulator of the dmF transcriptome (Fig. 3G). Sub-clustering of Schwann cells reveals that Pdgfb is expressed almost exclusively by peri-synaptic Schwann cells (Fig. 3D). This coupled with the co-localization of S100b, Pgp9.5 and Cd34 (Fig. 6B) indicates a signaling niche in the interstitial space of DSM muscle fascicles between peri-synaptic Schwann cells, neurons, detrusor smooth muscle, and fibroblasts (Fig. 6G).

Beyond the major function related elements of the bladder detailed above other cell types identified in this study included mesothelial, endothelial and immune cells. Interestingly, the mesothelial cluster shares the same marker genes with the mouse cl.11 of the Yu et al^48^ data set, defined as ‘neuron’. Given the high expression of the classical mesothelial marker *Msln* (mesothelin) and localization of a marker for this cluster, *Gpm6a*, to the traditional mesothelial region in the outermost layer of the bladder (Fig. 1F) we believe these are indeed mesothelial cells and not neurons. Subclustering of immune cells revealed 13 unique cell types (Fig. S5A&B) and when subclustering was performed with inclusion of a publicly available mouse PBMC dataset two myeloid populations (cDC2 and macrophages) were primarily bladder in origin indicating tissue residency (Fig.S5C). Markers for these cluster *Xcr1* (cDC2) and *Adgre1* (macrophages) have been known to mark tissue resident populations. Interestingly, certain myeloid populations displayed high layer specificity based on IMC and ST (Fig. S5D&E). Macrophage and MHCII+ monocytes were more likely to be located in the detrusor region while cDC2 was almost exclusive to the urothelium/lamina propria region (Figure S5E). Additionally, large groups of contiguous spots positive for the plasma cell marker *Jchain* were identified in ST data. Specifically these large clusters of contiguous spots were present in aged but not mature sections (contiguous spots Aged=462, Young=3, Fig.S6) which is in line with a recent report^49^ wherein tertiary lymphoid structures, which are highly populated with plasma cells, were more prevalent in aged bladder sections.

## DISCUSSION

We have generated a comprehensive cell atlas of the mouse urinary bladder by combining multiple techniques, both dissociative and those that maintain spatial organization, significantly improving upon past efforts to profile the mouse bladder at single cell resolution. In addition to greatly expanding the numbers of profiled individual cells and therefore gleaning insight into cell type sub-structure, we provide the first cell-type-specific transcriptomic view of the detrusor muscle cell, and identify age-specific dissociation differences which have confounded the interpretation of earlier single cell studies. Table S8 provides a summary of cell types we have defined with, where available, the cell ontology terminology from EMBL-EBI Cell Ontology. Bladder ontology terminology was noticeably sparse compared to other organs such as lung and heart indicating a lack of adequate bladder cell typing.

The limitations in dissociative techniques are evident in past studies of the bladder^5, 6, 50^ that nearly completely lack representation of the detrusor smooth muscle cell, a major component of this tissue. DSMs have strong intercellular attachments through detrusor muscle fibers and can be very large in size and therefore, similar to cardiomyocytes, are a challenge to isolate. With our combination of strategies focused on the detrusor layer and including single nucleus profiling we were able to generate 474 single nuclei and 342 single cell transcriptomes of DSM cells and validate expression localization to the detrusor layer through spatial methods.

Cells attached to or embedded in the extracellular matrix (ECM) can inhibit their recovery in dissociative techniques; we have previously seen this in the percent recovery of cells between the morula and late-stage blastocyst^51^ when ECM is first deposited, and in the efficiency of recovery of fibroblast in heavily desmoplastic tumors such as in pancreatic cancer^28^. This effect apparently confounded the interpretation of single cell data in a recent study concluding urothelial cells increase in numbers with age^32^. Here we show that the fibroblast-containing lamina propria has age-dependent biases when stripping urothelial from detrusor muscle layers, coming off more with the former in young mice and the latter in aged mice. This would lead to technical artifacts in the ratios of urothelial to fibroblast that were profiled in the Tabula Muris^32^. Instead, and in line with previous literature reporting increased fibrosis in aged bladders^52, 53^, our data suggest an increase in the proportion of fibroblasts and, specifically a distinctive increase in ECM-producing differentiated myofibroblast with age. Further, we provide evidence that TGFB1 is a contributor to this differentiation, a factor known to increase with aging^54^. Interestingly, the expression profile of these fully differentiated myofibroblasts include genes known to be associated with age-related bladder disorders such as neurogenic bladder dysfunction^55^. These differential dissociation outcomes of fibroblast with age emphasized the importance of not solely relying on one method of interrogation in such studies.

While we were careful in employing complementary techniques to capture a comprehensive view of cell types and their associated transcriptomes, our study still lacks in describing the biology of, and heterogeneity within, bladder neuronal innervation. Neuronal cell bodies are not present in the highly innervated bladder. We believe we capture some axonally-localized transcripts (e.g. *Npy*, *Slc17a7*), present in the bulk RNA-seq and spatial transcriptomics but absent in the single cell and nucleus data, but only slightly informs the rich biology in these structures. Future studies incorporating neuronal specific techniques such as retrograde tracing will better provide insight into the transcriptome, and therefore the biology, of these neurons.

In addition to the cataloging of cell types and states, single cell analysis can capture developmental transitions and provide insight into the molecular control of cell fate decisions^51^. With the numbers of individual cells profiled here we were able to reconstruct the differentiation dynamics found within the urothelial layer, from basal progenitors to the fully differentiated luminal umbrella cells, validating the pseudotime projections generated from the scRNAseq data with spatial transcriptomics mapped to the histology of this layer. This transcriptome dynamics aids in understanding the genetic regulatory network controlling the normal urothelial regenerative process but also how this may be co-opted in disease mechanisms. With respect to the latter it is interesting to note we find genes associated with a number of somatic mutations apparently selected for in the normal aging bladder^56^, and enriched in cancer lesions, are dynamically expressed through this process suggesting these pre-cancerous mutations are beginning to subvert the normal regenerative pathway within the urothelium.

The characterization of interstitial cells of Cajal (ICC) within the gastrointestinal (GI) tract as pacemakers for smooth muscle cell contraction via response to enteric motor neurotransmitters^57^ led to the search for similar cells controlling detrusor muscle function in the bladder. In the gut ICCs are localized between muscle fascicles and within the bladder a fibroblast subtype exists within the space between detrusor fascicles. However, as opposed to being ICC-like our data indicates this cell type to be closely related to other bladder fibroblasts and forms a continuous fibroblastic network extending from the suburothelium deep into the detrusor. The growth factor receptor Kit marks GI ICCs and its associated antibodies have been used with mixed results to define fibroblast-like ICCs within the bladder^23, 26^. Our data agrees with the absence of *Kit* expression in the fibroblast/interstitial cell compartment. In addition, our data indicates that other highly expressed pan-ICC markers identified within the gastrointestinal track (*Ano1*, *Gja1*, *Hprt*)^58^; are also either absent or extremely low in expression in bladder fibroblasts. These are not just markers for ICCs but functionally required for electrical activity in the mouse intestine ^59, 60^. While this does not rule out a role for fibroblasts in bladder volume control, the absence of an ICC-like cell within the bladder argues for fundamental cell-type differences in the control of smooth muscle contractility between the bladder and those lining the gastrointestinal tract.

Our ability to capture the detrusor muscle cell transcriptome, lacking in previously published single cell work, will provide further insight into bladder muscle control. Indeed, multiple neuronal-like genes, presumed contributors to the post-synaptic junction, are expressed within the detrusor smooth muscle cells and point towards new players at the neuromuscular junction. Intriguing in this regard is the DSM-specific expression of *Stum*, a molecule first described as mechanosensing in *Drosophila* proprioceptive neurons^44^ though with little functional information yet described in mammals. Evidence for a signaling niche that involves DSM, fibroblasts and neurons has been suggested^61^ and our spatial data supports this and adds peri-synaptic Schwann cells to the mix. Importantly, with the cell-type-specific transcriptomes of three of these components now in-hand – the neuronal transcriptomes were not captured in this study - insights into this signaling niche can be gained, an example of which being the aforementioned PDGBB induced differentiation of fibroblasts within the detrusor region.

The mouse is an essential model to understand the biology of the human bladder as it remains challenging to establish a human-based *in vitro* model that can fully replicate the integrated physiology of the bladder (*e.g.* filling, voiding, innervation). As mouse gene knock-out efforts continue^62^, having comprehensive tissue atlases such as we have generated here, will guide subsequent phenotyping efforts, which are currently underway for *Stum* and *Gm28729*. As a model of human biology, it is essential to define the similarities and differences between the mouse and human. The expectation is that much of the biology will be conserved but identifying where this varies will further inform the biology of both systems. We noted an intriguing species difference in DSM-specific *Gm28729*, a 409 amino acid protein coding gene in the mouse that remains without functional annotation, with a high bladder-specificity score, conserved across at least 350 million years of time, yet apparently recently lost in humans and Denisovans (Fig. S3). While functional characterization awaits, one may speculate on an association between this mutation and bladder-specific features of the hominins. In this regard, it is noteworthy that an association between bipedal gait, a hominin-specific feature, and urinary bladder control has been identified^63, 64^. Future comprehensive cell-type-specific gene expression comparisons between the mouse and human bladder will further inform the similarities and differences in this organ.

Overall, this study represents the most comprehensive atlas of the mouse bladder to date. This resource provides a foundation of bladder cell types from which researchers can elucidate aspects of bladder physiology and pathobiology. This data is available in multiple formats from the raw sequence trace files and associated count matrices to analyzed and interpreted outcomes (https://singlecell.jax.org/bladder). Importantly, we show true tissue atlases require a combination of tissue processing and analytical techniques, with age of tissue-dependent considerations, providing important context for the rapidly growing cell atlas construction community.

## DATA ACCESS

All raw and processed sequencing data generated in this study including scRNAseq, snRNAseq, Visium ST, and bulk RNA seq have been submitted to the NCBI BioProject database (http://www.ncbi.nlm.nih.gov/bioproject) under accession number GSE180128. Publicly available datasets can be downloaded from Tabula Muris Consortium (https://tabula-muris.ds.czbiohub.org/), Mouse Cell Atlas (Microwell-Seq) (http://bis.zju.edu.cn/MCA/) and 10X (https://www.10xgenomics.com/resources/datasets) websites. To provide data accessibility and allow use as a resource, we made our processed scRNAseq, snRNAseq and Visium Spatial Transcriptomics data available at https://singlecell.jax.org/datasets.

## Supporting information

TableS1_Details_of_Datasets

TableS2_DE_Bulk_vs_Initial_Datsets

TableS3_scRNA_seq+Bulk_Counts

TableS4_IMC_Panel

TableS5_Myofibroblast_DE

TableS6_Smooth_Muscle_Markers

TableS7_Visium_Neuronal_Spot_DE

TableS8_Marker_Genes_&_Cell_Ontology

TableS9_Luminal_vs_Basal_DE_Visium+scRNAseq

## ACKNOWLEDGEMENTS

FUNDING: NIH/NIA: K76AG054777 (PPS), R01AG058814 (PPS), Travelers Chair in Geriatrics and Gerontology (GAK). The Jackson Laboratory startup funds (PR).

## AUTHOR CONTRIBUTIONS

Conceptualization – P.R., P.P.S. Methodology – D.B., I.M.A., S.S., A.A. Software – W.F.F., D.B. Formal Analysis – D.B. Investigations – D.B., D.L., A.A., S.S. Resources – P.P.S., P.R. Data Curations – D.B., W.F.F. Writing Original Draft – D.B., I.M.A., P.P.S., P.R. Writing Review & Editing – G.A.K., W.F.F, D.B., I.M.A., P.P.S., P.R. Visualization – C.C.H., D.B. Supervision – P.R., P.P.S., G.A.K. Project Administration - P.R., P.P.S., G.A.K. Funding Acquisition - P.R., P.P.S.

## STAR METHODS

### Bladder Harvesting

All animal procedures were conducted according to protocols approved by the University of Connecticut Health Center Animal Care Committee and the JAX Animal Care and Use Committee. Animals (C5BL6/J mouse) were sacrificed by CO_2_ euthanasia and bladders harvested into cold PBS or media, depending on the dissociation protocol, and individually minced using microscissors prior to enzymatic dissociation.

### Dissociation Methods for scRNAseq

Three different bladder tissue dissociation protocols were used in this study (described below) to achieve optimal cell type representation. Dissociation Protocol 1 was adapted from Mora-Bau et al.^66^ as a general bladder dissociation technique. Dissociation Protocol 2 was developed as a modified Dissociation protocol 2 in order to increase cell viability prior to sorting. Dissociation Protocol 3 was adapted from Hristov et al.^24^ to specifically dissociate cells from the detrusor smooth muscle layer.

#### Dissociation Protocol 1

Bladders were collected in cold PBS and, after mincing, each bladder transferred to 1 mL of freshly prepared digestion solution (0.06mg/mL Liberase TM (Roche), 0.125mg/mL DNAse I (Roche) in PBS), and incubated for 1 hour at 37⁰C in a water bath with vigorous shaking at 15 min intervals until a glassy appearance was reached. Reactions were stopped by addition of cold FACS buffer (2% FBS, 0.2 mM EDTA in PBS). Cells were filtered using a 100 μm filter (Corning), collected by centrifugation and resuspended in FACS buffer (2% FBS, 5mM EDTA in PBS).

#### Dissociation Protocol 2

Bladders were collected in cold Ham’s F10 medium (Gibco) and, after mincing, each bladder transferred to 2 mL of freshly prepared digestion solution (0.06mg/mL Liberase TM (Roche), 0.125mg/mL DNAse I (Roche) in DMEM/F-12 medium (Gibco)), and incubated for 15 min at 37⁰C in a water bath with gentle inversion every 5 min. After a final trituration larger material/tissue was allowed to settle for 1 min and cells in suspension collected and placed into cold FACS buffer on ice. This procedure was repeated two more times on the remaining tissue. The combined cells were filtered using a 100 μm filter (Corning), collected by centrifugation, RBCs removed by ACK lysis (Gibco) then washed with FACS buffer. Cells were stained with DAPI (Sigma) and CalceinAM (Life Technologies) and live cells collected by FACS.

#### Dissociation Protocol 3

Bladders were collected in cold PBS and using forceps, mucosa and detrusor were separated and minced separately. After mincing, mucosa tissue was processed according to Dissociation 2 up until ACK lysis. Detrusor tissue was transferred to 2 mL of a freshly prepared papain solution (1 mg/mL BSA (Sigma), 1 mg/mL papain (Stem Cell Technologies), 1 mg/ml DTT (Sigma) in Ca^2+^-free PBS), and incubated at 37⁰C in a water bath for 25 min with gentle inversion every 5 min. Detrusor tissue was washed 2x with Ca^2+^-free PBS and transferred to 2 ml of freshly prepared collagenase solution (1 mg/mL BSA, 1 mg/mL collagenase II (Sigma), 100 μM CaCl2 (Teknova) in Ca^2+^-free PBS), incubated 10-15 min at 37⁰C in a water bath with gentle inversion every 5 min. Detrusor tissue was then gently triturated and cells in suspension collected and placed into FACS buffer, filtered using a 100μm filter and collected by centrifugation. Detrusor and mucosa samples were counted by Countess II (Thermo Fisher) and pooled in equal amounts for library preparation.

##### Flow Cytometry and Sorting

C5BL6/J mouse bladders were dissociated into single cell suspensions and stained with Dapi and CalceinAM (Dissociation 1/2 above). Cells were stained with Dapi (Sigma) and CalceinAM (Life Technologies) according to manufacturers instruction prior to sorting. Live cells were sorted using FACSAriaTM Fusion (BD Biosciences). After exclusion of debris and doublets, Dapi- and CalceinAM+ cells were sorted as live viable fraction (50,000-100,000 cells per sample) and collected in 2% FBS/PBS + EDTA (FACS buffer). Sorted viable cells were then washed and resuspended with 0.01% BSA in PBS and assessed for viability using trypan blue staining for subsequent scRNAseq experiments. Flow cytometry analysis for assessment of mesenchymal:urothelial ratio in age and young mouse bladders was performed by dissociating cells by Dissociation 1 and collecting cells in FACS buffer and blocked with anti-mouse CD16/CD32 Fc Block ( BioLegend, clone 2.4G2, 1:50) for 15 minutes . Staining of cell suspensions was performed with CD326 APC (urothelial marker) (eBioscience, clone G8.8) and V-CAM1 FITC (mesenchymal marker) (Thermo Fisher clone M/K-2). Prior to sorting, Dapi was added to cell suspension (Sigma). After debris, doublet and dead cell exclusion, mesenchymal:urothelial ratio was determined as the number of V-CAM1+ cells / CD326+ cells. For assessment of layer specific age-related changes in cell proportions 22 month and 4 month mice were sacrificed and bladders were removed. Using forceps, mucosa and detrusor were separated and dissociated separately by Dissociation 2 for mucosa and Dissociation 3 for detrusor. After dissociation cells were resuspended in FACS buffer and stained with CD34 AlexaFluor 647 (RAM34 eBioscience) (fibroblast marker) for 30 minutes on ice Cells were then washed by centrifugation, fixed with 4% PFA and washed again by centrifugation again prior to flow cytometry. Cells were analyzed using BD LSRII (BD Biosciences) and fibroblast percentage for each sample determined as number of fibroblasts / total number of cells after debris and doublet exclusion. All post hoc flow cytometry analysis was performed withFlowjo (version 10).

### Nuclei Preparation

Fresh bladders were mechanically stripped to separate detrusor and mucosal layers. Each layer was then minced and flash frozen on dry ice. Tissue was placed in 50 μl of ice cold nuclei EZ lysis buffer (Sigma) and ground by mortar and pestle for 5 min (SP Bel-Art). Sample were then centrifuged 500g for 5 min at 4⁰C. Supernatant was removed and 50 μl of fresh lysis buffer was added and cells were incubated for 5 min after which 50 μl of PBS with 0.04% BSA and Ambion RNAse inhibitor (Invitrogen) were added. Samples were then washed with PBS with 0.04% BSA and filtered with a 40 μm filter followed by an additional wash and filtration with a 5 μm filter. Nuclei were then counted with a Countess II (ThermoFisher) and equal numbers of nuclei from each matched detrusor and mucosal preparations pooled prior to loading on 10x chromium chip.

### Single-Cell/nuclei Capture, Library Preparation, and RNA-seq

Prepared cells or nuclei in PBS containing 0.01% BSA were quantified on a Countess II (Thermo Fisher), and up to 12,000 cells/nuclei were loaded per channel on a Chromium microfluidic chip (10x Genomics). Single- cell/nuclei capture, barcoding, and library preparation were performed using Chromium version 1, 2, or 3 chemistries according to the manufacturer’s protocols (#CG00103 10x Genomics). cDNA and library quality were verified on an Agilent 4200 TapeStation and libraries quantified by KAPA qPCR before sequencing (HiSeq4000/Novaseq, Illumina) targeting an average depth of 50,000 reads per cell.

### Single-Cell Data Processing, Quality Control, and Analysis

Illumina base call (BCL) files were converted to FASTQ files using bcl2fastq (Illumina, version 2.16.0.10). CellRanger 3.1.0 (10x Genomics) was used to align FASTQs to the mm10-3.0.0 reference (Ensembl build GRCm38.84) and produce a digital gene-cell counts matrix. Publicly available counts matrices^5, 6^ were downloaded from Tabula Muris and Microwell Cell Atlas websites respectively. Subsequent data processing was performed in python utilizing the Scanpy 1.4.6 package^64^. Gene-cell matrices were filtered to remove cells with fewer than 500 transcripts and genes with fewer than 3 counts and present in more than 3 cells. Individual gene-cell matrices were then normalized such that the number of unique molecular identifiers (UMI) in each cell is equal to the median UMI count across the data set and log transformed. Cells with over 30% mitochondrial transcripts in non-detrusor enriched samples and cells with over 20% hemoglobin transcripts were filtered from downstream analysis.

Samples were then aggregated and the top 2,000 genes with the highest variance across the aggregated dataset were identified based on their mean expression in the population and dispersion. Highly variable genes were used as input to dimensionality reduction. Batch corrected dimensionality reduction was performed as follows: PCA embeddings were generated and corrected for library preparation and dissociation technique using Harmony^68^. Corrected PCA embeddings were used to generate nearest neighbor graph using BBKNN^69^ and used for dimensionality reduction via UMAP. The resultant UMAP embeddings were clustered via scanpy built in Leiden community detection algorithm to produce labeled cell clusters. Marker genes were identified using a one-versus-rest strategy which determines marker genes by area under a receiver operating characteristic curve (AUROC) analysis for all genes that are greater than twofold expressed in a cluster compared with all other cells. Genes with greater than 85% AUROC were defined as markers specific to the cell type. All differential gene expression analysis was performed using edgeR (version 3.28.1)^70^. Subcluster analysis was performed by subsetting the global dataset according to cell type of interest and performing dimensionality reduction, clustering and marker identification as above on the subset.

Processed scRNA/snRNA datasets were aggregated to create pseudobulk datasets by averaging transcriptomes across all cells for comparison to processed averaged bulk RNA datasets generated from dissociated and whole bladders. Mapping of neuronal transcripts in detrusor smooth muscle was confirmed by visual inspection the snRNA seq and bulk whole bladder samples in IGV 2.3.32^71^.

Pseudotime analysis of urothelial clusters was performed using PAGA^72^ . Briefly, relevant cell clusters were extracted and subclustered by using the neighborhood graph from above followed by dimensionality reduction via with force-directed graph drawing (Fruchterman Reingold) and Louvain clustering. Urothelial gene trajectory analysis was performed by binning cells (n=8) at uniformly distributed intervals across pseudotime and averaging gene expression across cells in each bin to generate a pseudotime-gene expression matrix. The gradient of the pseudotime- gene expression matrix was then used as input into UMAP to generate trajectory embeddings.

To quantify the percent of terminally differentiated suFs with respect to age, the suFs were extracted from batch matched scRNAseq datasets containing old and young/mature samples (WB1+WB2, WB3+WB4+WB5). For each set of batch matched samples, gene expression matrices of known myofibroblast differentiation genes were extracted and fit to a gaussian mixture (n_components=2) to classify suFs as terminally of partially differentiated. Significance was determined by Fisher’s Exact Test.

For aging transcriptome analysis, genes found to be significantly differentially expressed between mature and aged mice based on Kamei et. al.^65^ were identified on GEO (GSE100219) and downloaded for plotting within scRNAseq datasets. Specificity indices were generated by downloading Mouse ENCODE transcriptome data from the NCBI Gene website (https://www.ncbi.nlm.nih.gov/gene/) and bladder specificity determined by bladder expression / total expression in all other tissues. Regulator analysis of fibroblasts was performed with IPA (QIAGEN Inc., https://www.qiagenbioinformatics.com/products/ingenuity-pathway-analysis) using edgeR DE output between suF and dmF as input into IPA. To co-cluster bladder immune cells and PBMC’s, a publicly available C57BL/6 peripheral blood mononuclear cell (PBMC) scRNA-seq gene counts matrix (5’ Gene Expression V1) was downloaded from the 10X website (www.10xgenomics.com/resources/datasets) as a counts matrix. The public PBMC counts matrix was concatenated with acounts matrix generated from immune cell subclusters from our in house datasets. The combined dataset was processed according to filtering and batch corrections metrics stated above with additional batch correction for cell origin (PBMC, bladder).

### Cross species comparison *GM28729*

Predicted protein sequence was obtained from the ProteomicsDB (https://www.proteomicsdb.org/), compared to other organisms with NCBI BLAST (https://blast.ncbi.nlm.nih.gov/Blast.cgi) and visualized with MView (https://www.ebi.ac.uk/Tools/msa/mview/).

### Imaging Mass Cytometry (IMC)

15 antibodies were used for IMC, a full description of these can be found in Table S4. Antibodies selected for the panel were first validated via immunofluorescence and subsequently metal-conjugated using the Maxpar X8 Multimetal labeling kit (Fluidigm). PFA-fixed OCT embedded mouse bladder tissues were cut into 5 μm sections and mounted on slides. After blocking in a buffer containing 10% BSA, slides were incubated overnight at 4°C with a cocktail of metal-conjugated IMC-validated primary antibodies. The following day, slides were washed twice in PBS and counterstained with iridium intercalator (Fluidigm) (0.25 μmol/L) for 5 min at RT, to visualize the DNA. After a final wash in ddH_2_0, the slides were air-dried for 20 min. The slides were then loaded on the Hyperion/Helios imaging mass cytometer (Fluidigm). Regions of interest were selected using the acquisition software and ablated by the Hyperion and metals measured by the Helios. The resulting images were exported as 16-bit .tiff files using the MCDViewer software (Fluidigm) and analyzed using the open source Histocat++ 1.76 toolbox and ImageJ 1.53. Immune cell type localizations from IMC were determined in ImageJ by manually separating detrusor and urothelium/lamina propria regions into separate images and using automated counting for cell specific marker positive objects in each region.

### Bulk RNA Sequencing

12-month old male and female mouse bladders were bisected and one half dissociated into single cells using Dissociation 2 while the other half was directly homogenized without single cell dissociation. RNA was extracted using RNEasy kit (Qiagen). Library preparation was performed using TruSeq RNA Library Prep Kit v2 (Illumina). Libraries were sequenced on an illumina Hiseq 4000 at a depth of 80 million reads per library. Data was processed with bcl2fastq2 (version 2.16.0.10) to generate FASTQ files. FASTQs were aligned via STAR 2.7.3a to the mm10-3.0.0 mouse reference genome used for scRNAseq analysis and a transcript counts matrix generated with subread 1.5.2 featureCounts. Counts were normalized by TPM. Differential expression between groups was performed with edgeR (version 3.28.1).

### Spatial Transcriptomics

Spatial transcriptomics was performed according to the Visium Spatial Gene Expression Solution (#CG000239 10x Genomics). First, the optimal length of time for permeabilization was determined to be 20 min from using a Visium spatial tissue optimization slide (10x Genomics) testing times of 3, 6, 12, 18, 24, and 30min on 10 μm thick sections of an OCT-embedded, unfixed mouse bladder. To generate the spatial transcriptomics data four 12- month and four 22-month old bladders were embedded in OCT without fixation, in sets of two each, and snap frozen for a total of 8 blocks. From this, one 10 μm thick section from each block (each containing two bladders) was carefully positioned across one 6.5 mm^2^ capture area on the Visium Spatial Gene Expression slide with four areas per slide for a total of two slides. Slides were stained with H&E, imaged on a Phenix High Content Imaging system (Perkin Elmer) with a 20x objective prior to a 20 min permeabilization. Sequencing was performed on an Illumina Novaseq with 300 million reads per library. Image alignment was performed with Space Ranger V1.1 and data analysis was performed with scanpy^67^. Cluster generation was performed in the same manner as scRNAseq data with the exception of dimensionality reduction feature selection in which genes identified as cell type markers in scRNAseq data were used as input. For more specific localization of particular subclusters identified in scRNAseq data fibroblast the gene list for dimensionality reduction was adjusted to include subcluster specific markers. To identify neuronal signatures in ST data, spots with *Npy* or *Slc17a7* expression > 0 were annotated as neuronal and DE analysis with edgeR was performed between neuronal and all other spots. For immune and Schwann cell localizations by Visium ST, spots were annotated as containing immune or Schwann cells based on thresholding expression of markers genes identified in scRNAseq data and classified as detrusor or uro/lp based on regional position using grayscale H&E images. For basal vs. luminal urothelium comparison, grayscale H&E images were used to manually annotate fiducials in Loupe Browser (version 4.0.0) as basal or luminal based on known locations of these cell types relative to the bladder lumen. Barcodes of basal and luminal annotated spots were exported and used to subset and annotate ST expression matrix for DE analysis. For comparison of tertiary lymphoid structures (TLS) between mature and aged ST samples, a plasma cell marker (*Jchain*) known to localize to TLS was used to identify spots *Jchain* expressing spots with neighboring spots (n>2) also expressing *Jchain*.

### Tissue culture

Fibroblasts were derived from 12-month-old mice by stripping the mucosal layer from the bladders and processing this into a single cell suspension according to Dissociation 3 protocol described above. Cells were then stained with AlexaFluor 647 CD34 (RAM34 eBioscience) and calcein AM (Life Technologies) for 30 min and propidium iodide (BD Pharmingen) for 10 min. Cells were washed with FACS buffer and sorted on a FACSAria^TM^ Fusion (BD Biosciences). Viable CD34+ cells were collected in media (10% FBS (Gibco), MEM non-essential amino acids, 100 U/mL primocin (Invitrogen) in Advanced DMEM/F12 (Gibco) and cultured on Matrigel-coated (Corning) 24-well plates (Corninng) at 50,000 cells/well with 500 μl of media/well. The following day, when cells were confluent, media was exchanged with media alone, media with 20 uM AG370 (Enzo Life Sciences) and 10 ng/ml TGB1 (LS Bio), or media with 3 uM A83-01 (Tocris) and 10 ng/ml PDGFBB (Stem Cell Technologies). Cultures were maintained for 6 days with media changes occurring every other day. At day 6 RNA was isolated utilizing Arcturus PicoPure RNA Isolation Kit and qPCR reactions were prepared using SuperScript III Platinum Kit with SYBR Green One Step qRT PCR with the following primers from IDT: Penk: F-ACACAACTTCACTAATCCAGGTG R- GAAGCCTCCGTACCGTTTCAT Acta2: F- GTCCCAGACATCAGGGAGTAA R- TCGGATACTTCAGCGTCAGGA Dlk1: F- CCCAGGTGAGCTTCGAGTG R- GGAGAGGGGTACTCTTGTTGAG Gpx3: F- CCTTTTAAGCAGTATGCAGGCA R- CAAGCCAAATGGCCCAAGTT Cxcl14: F- GAAGATGGTTATCGTCACCACC R- CGTTCCAGGCATTGTACCACT. IDT Ready-Made Gapdh F/R (51-01-07-12/13) was used for normalization.

#### Immunofluorescence on Frozen Sections

Bladder sections for immunofluorescence were prepared in the same manner as for IMC. Slides were permeabilized and block (30 minutes with 5% normal goat serum and 0.3% Triton X-100 in PBS) and incubated with primary antibodies in PBS with 0.03% Triton (Sigma) and 2% serum at 4°C overnight. The following day slides were washed with ice cold PBS and incubated with secondary antibody for 2 hours at RT. Slides were then washed with PBS + Hoechst (Invitrogen) and coverslips mounted (Prolong Diamond Antifade Mountant). Immunofluorescence images were acquired with a Zeiss Axiovert microscope at 20X. Primary antibodies used for immunofluorescence were PGP9.5 (PA5-85273, ThermoFisher), PGP9.5 (MA5-12371, ThermoFisher), DLK1 (LS-C746855-50, LS Bio), S100B (AMAb91038, Atlas Antibodies) and CD34 (14-0341-82, ThermoFisher). Secondary antibodies used were Goat Anti-Rat AlexaFluor 647 (A-21247 ThermoFisher), Goat Anti-Mouse AlexaFluor 568 (A-11004 ThermoFisher), Goat Anti-Mouse Goat AlexaFluor 488 (A-11001 ThermoFisher), Anti-Rabbit AlexaFluor 568 (A-11011 ThermoFisher), and Goat Anti-Rabbit AlexaFluor 488 (A-11008 ThermoFisher).

## Supplemental Information titles and legends

There are seven supplemental figures and nine supplemental tables as follows:

Table S1. Animals Used and Associated Datasets Generated

Table S2. Differential Expression Gene List Between Bulk and Single Cell RNA-seq

Table S3. scRNAseq and Bulk RNAseq Counts

Table S4. Details of Antibodies in IMC Panel

Table S5. Differential Expression of Myofibroblasts

Table S6. Markers of Smooth Muscle Cell Sub-types

Table S7. Neuronal Transcript Detection in Visium Data. Worksheet1 = Npy-associated genes, Worksheet2 = Slc17a7-associated genes

Table S8. Marker Genes Associated with Cell Ontologies

Table S9. Luminal versus Basal Urothelium Differential Expression. Worksheet1 = Visium, Worksheet2 = scRNAseq

---

**Figure S1.**
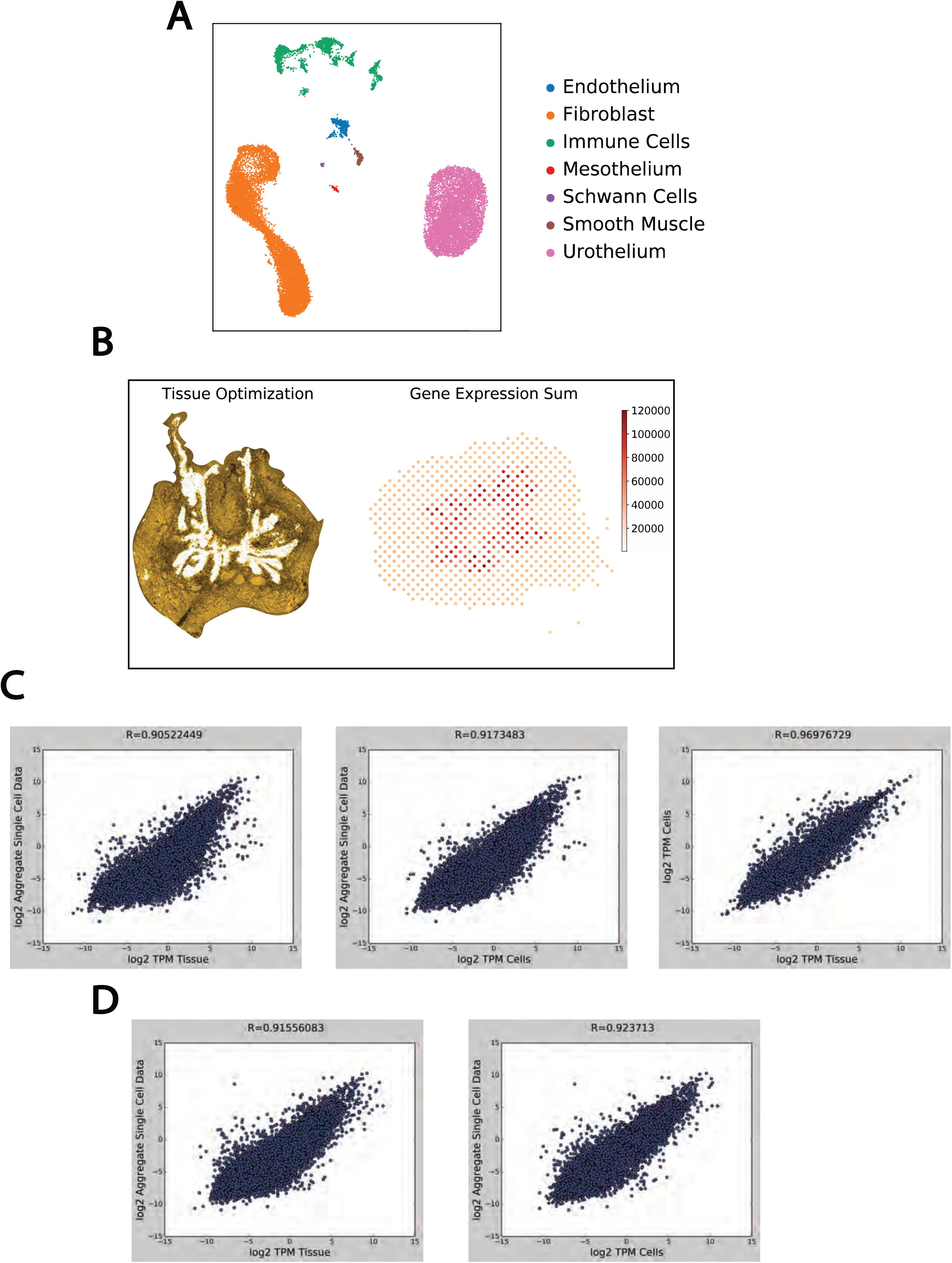
Initial Dataset Generation QC. A) UMAP plot color coded for cell types identified in initial datasets (see Table 1). B) Example image from Visium Tissue Optimization Protocol (left) indicates high RNA content (TRITC) in urothelial region and example UMI counts per spot (Gene Expression Sum) from processed ST section (right). C) Correlation plot of gene expression between pseudo-bulk initial scRNA seq datasets and bulk RNA sequencing (Whole Tissue + Dissociated Cells). D) Correlation plot of gene expression between pseudo-bulk complete scRNA seq datasets and bulk RNA sequencing (Whole Tissue + Dissociated Cells).

**Figure S2.**
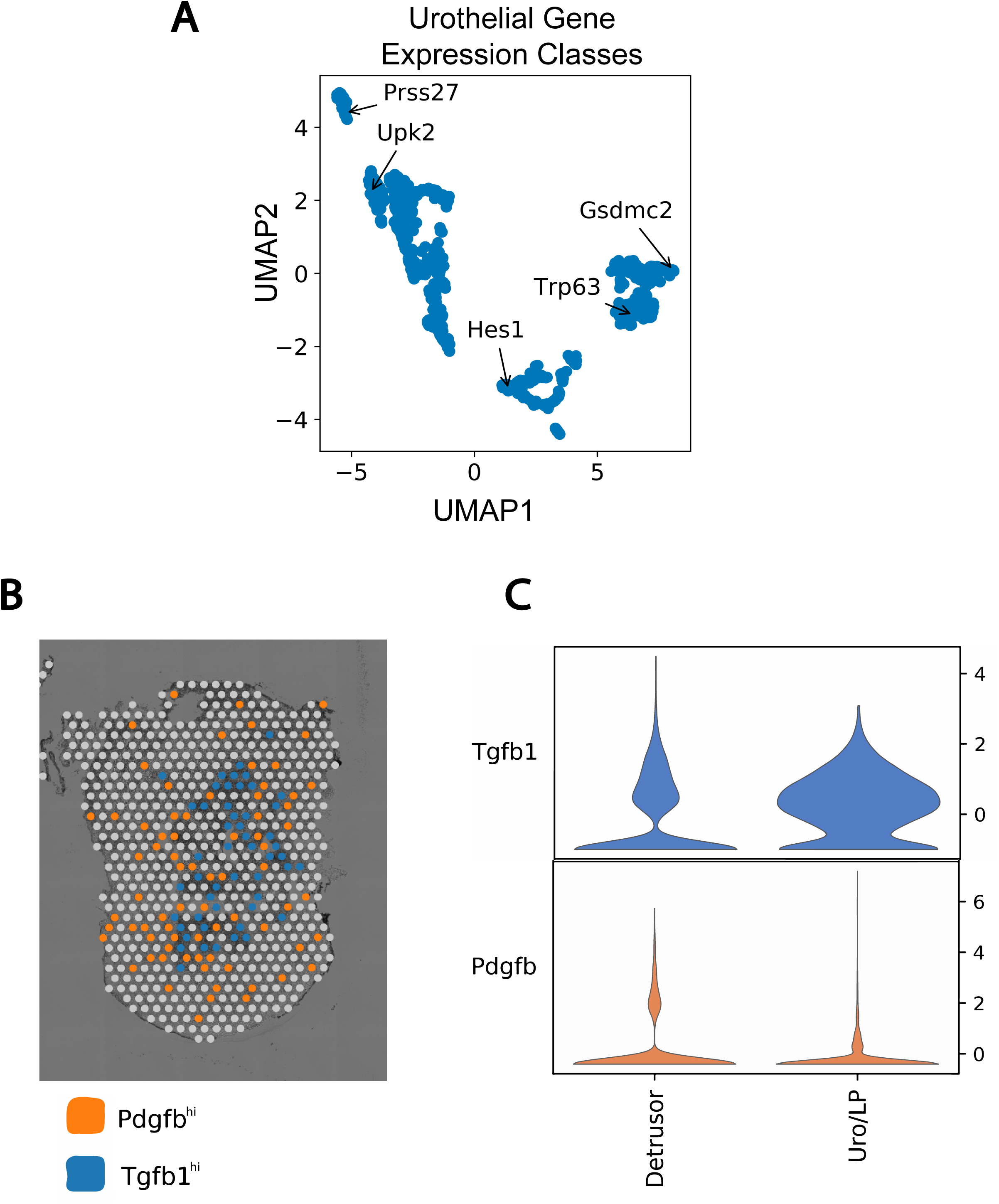
Urothelial Cell Pseudotime Analysis and *Pdgfb* and *Tgfb1* Localization. A) Gene expression across pseudotime was binned to generate a distinct pseudotime trajectory profile for each gene. 2D UMAP dimensionality reduction was applied to these profiles to identify groups of genes with similar dynamics. Five groups were observed corresponding to sets of early basal (*Gsdmc2*), basal (*Trp63*), intermediate (*Hes1*), luminal (*Upk2*) and late luminal (*Prss27*). B) Example ST plot of *Pdgfb* and *Tgfb1* high expressing spots. Higher expression of *Tgfb1* was observed within the urothelial region while *Pdgfb* was more evenly distributed across the whole bladder. C) Violin plot of *Tgfb1* and *Pdgfb* expression by region across all Visium datasets. Urothelial and lamina propria spots were binned together and *Tgfb1* and *Pdgfb* expression compared to detrusor region.

**Figure S3.**
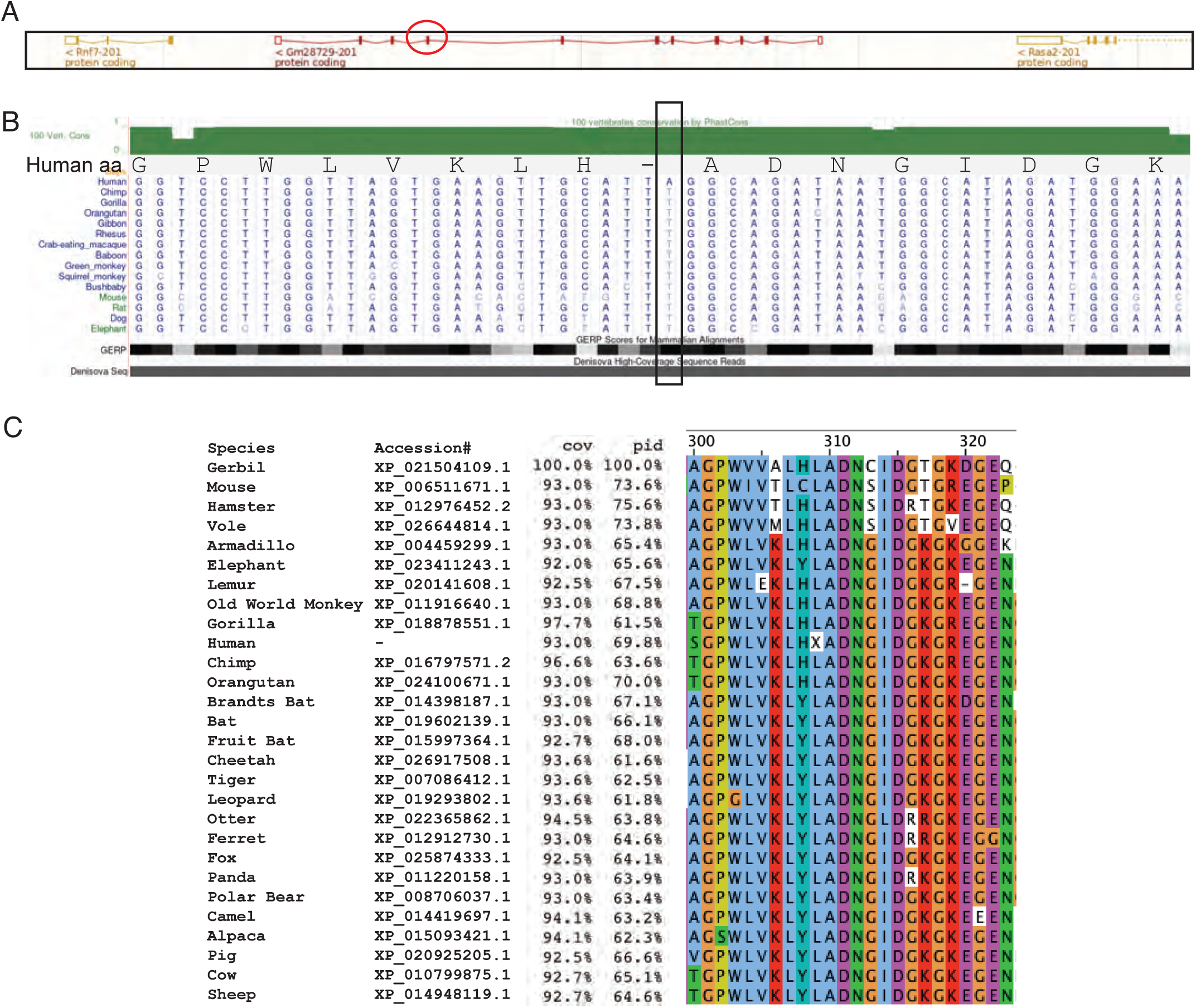
Protein alignment of GM28729 across Eutherians. A) A screen grab of the genomic region encompassing Gm28729 from the Ensembl genome browser from the mouse (GRCm39) chr9:96350407-96433904 inclusive of neighboring genes *Rnf7* and *Rasa2*. Exon 8, in which the homologous human sequence contains a nonsense mutation, is circled in red. B) 50 base region of human exon 8 sequence (hg19; chr3:141,432,169-141,432,218) encompassing the nonsense mutation (boxed) and visualized in the UCSC Genome Browser. PhastCons and genomic evolutionary rate profiling (GERP) tracks are included for a measure of sequence conservation. The PhastCons and GERP scores at the position of the nonsense mutation are 1 and 4.52, respectively, indicating high conservation score across vertebrates and mammals. A track for Denisova sequencing reads (Display mode – dense; representative of 29 reads spanning the site of interest) indicates identical sequence to that of human. An additional 14 non-human mammalian sequences, including three great apes, indicate the nonsense mutation is restricted to human and Denisovan genomes. Amino acid translation (Human aa) of the human sequence is shown with the stop codon (TAG) introduced by the nonsense mutation indicated by ‘-‘. All other extant mammalian sequences contain a TTG codon encoding a leucine (L) at this position. C) Protein sequence alignment scores of Gm28729 from 28 mammalian species generated from the indicated XP_ accessions. As the human gene (LINC02618; ENSG00000242104) is annotated as non-coding, the human protein sequence was manually generated from this gene guided by alignment to the mouse. The minimum, maximum, and average protein sequence length of the 28 sequences was 406, 462, and 415 amino acids, respectively. The alignment percent coverage (cov) with respect to the gerbil sequence and the corresponding percent amino acid identity (pid) are indicated. Protein sequence alignment is shown (right) for the region around the nonsense mutation which is indicated by an ‘X’ at position 309 in the human sequence.

**Figure S4.**
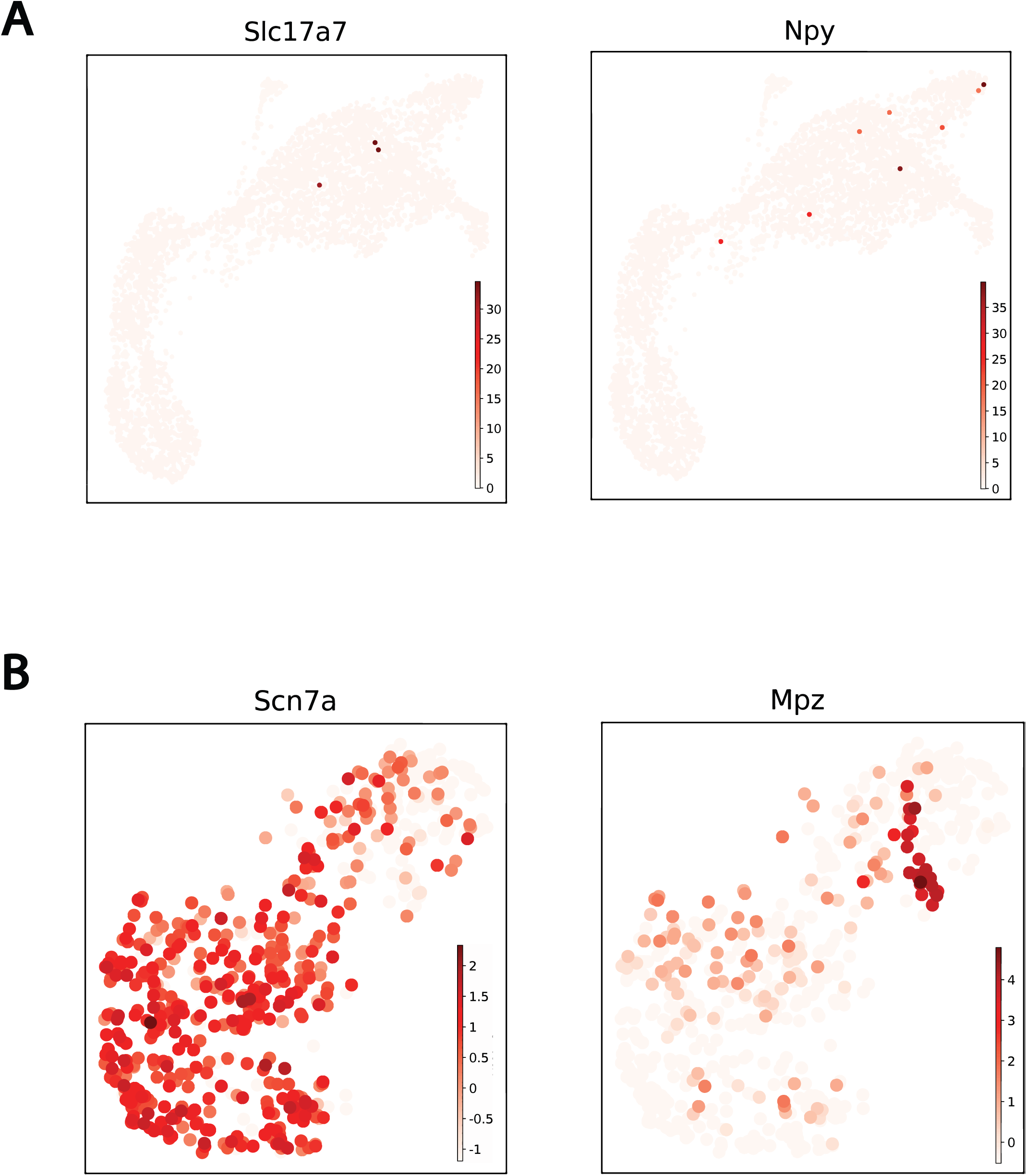
Identification of neuronal and Schwann cell marker genes. A) UMAP projection of markers identified as likely neuronal markers derived from Visium ST. Few *Slc17a7*+ (3) and *Npy*+ (8) spots were found. B) Marker genes used to identify Schwann cell subtypes upon subclustering. *Scn7a* and *Mpz* are well known perisynaptic and myelinating Schwann cell markers.

**Figure S5.**
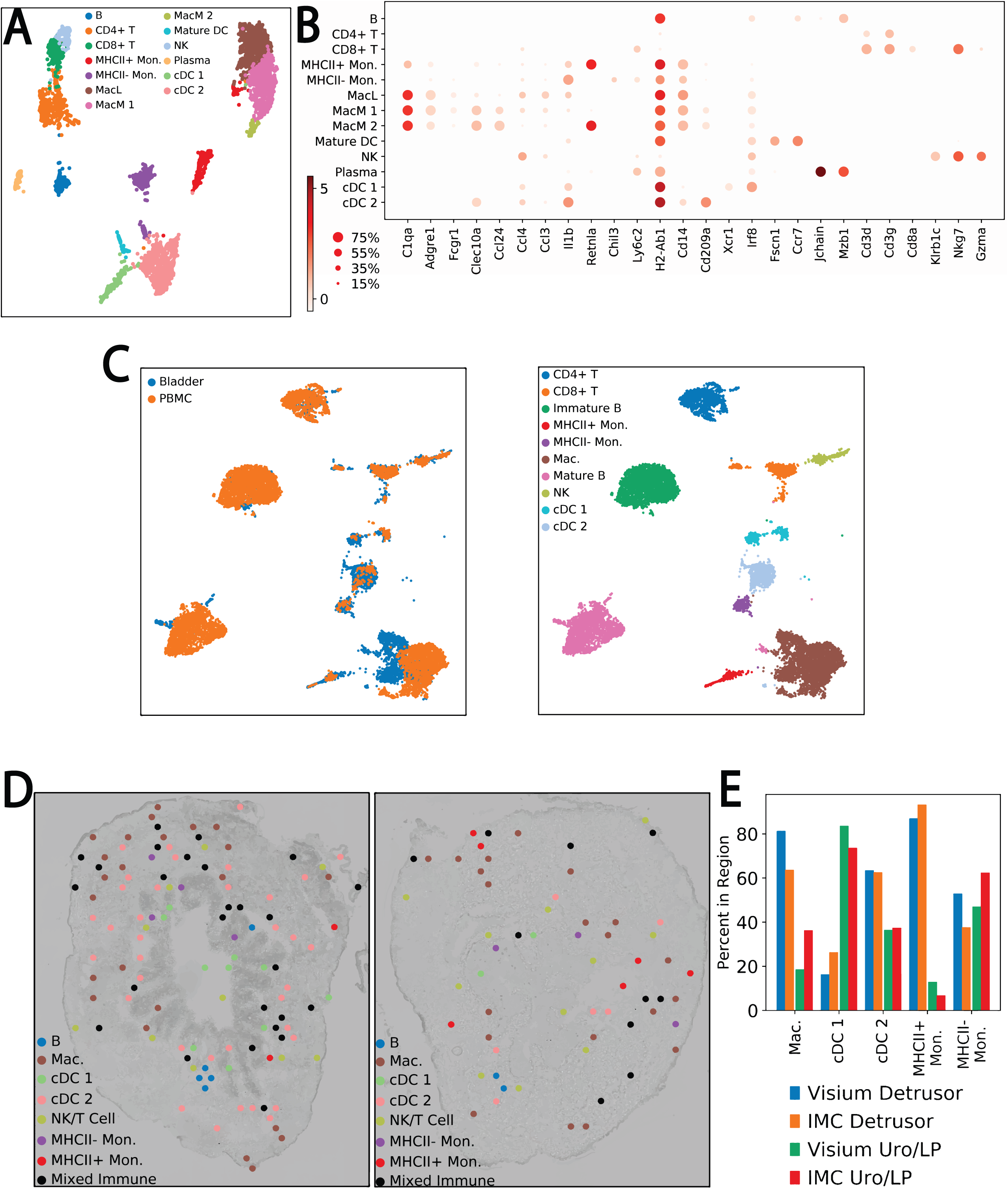
Immune Cell Composition of the Urinary Bladder. A) Subclustering of immune cell types results in 13 distinct clusters with 8 myeloid and 5 lymphoid subtypes. B) Heatmap of identified immune cell subcluster marker genes used to annotate clusters. C/D) Spatial localization of immune cell types based on Visium and IMC data. Visium and IMC images were split into detrusor and urothelium/lamina propria regions and the proportion of each immune cell type cluster was calculated. Majority detrusor groups included macrophages, cDC1, and MHCII+ monocyte. cDC2 was primarily Uro/LP with MHCII- monocyte displaying mixed localization. E) Immune cells were clustered with mouse PBMC data in order to identify potential bladder resident cell types based on overrepresentation of bladder origin cells vs PBMC origin. Myeloid cell types comprised the bladder origin clusters including a bladder origin macrophage subcluster.

**Figure S6.**
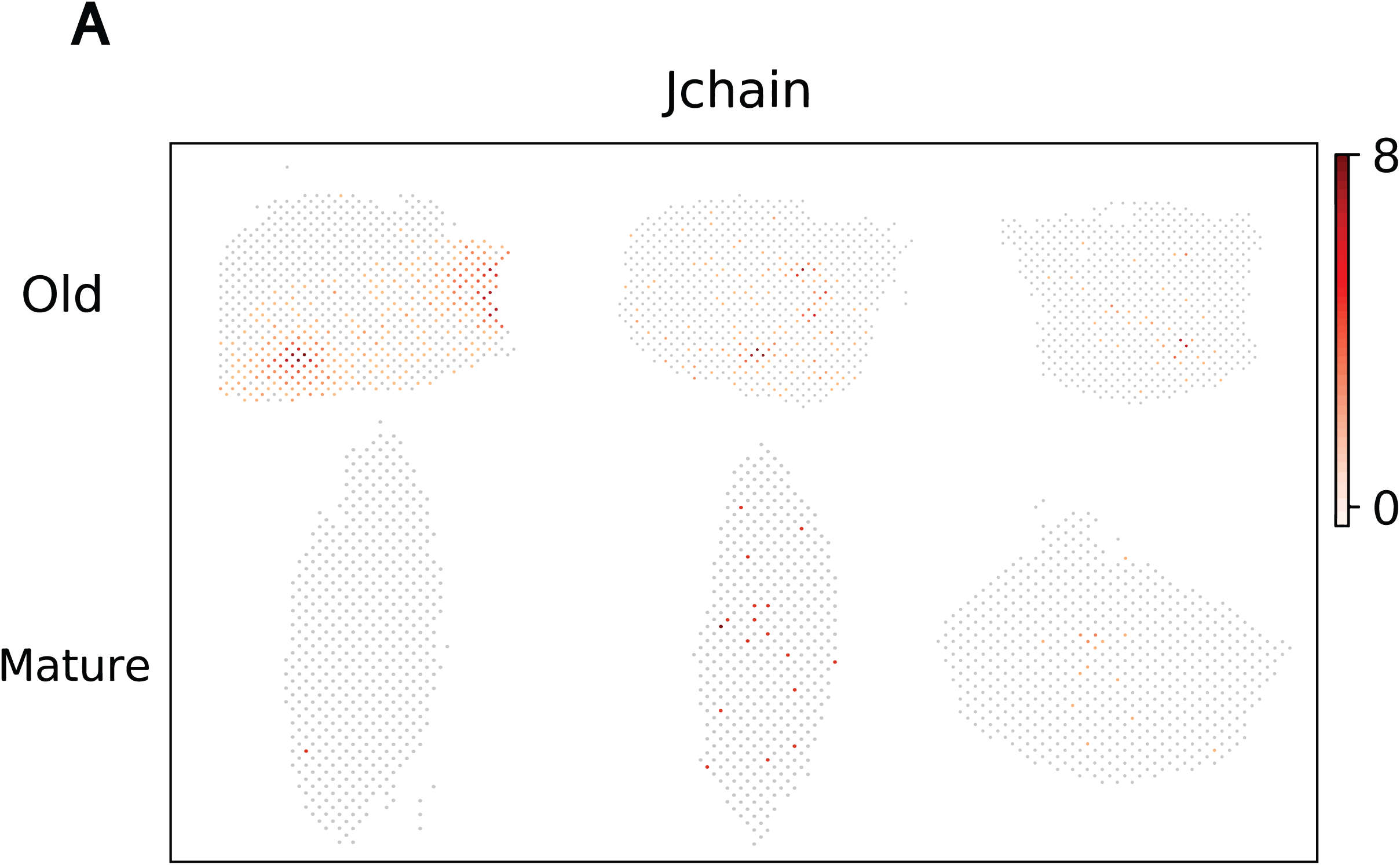
Plasma cell identification. A) Tertiary Lymphoid Structures in Aged and Mature ST Sections. *Jchain+* ST spots with neighboring *Jchain+* spots (n>2) were used to denote the potential presence of tertiary lymphoid structures in aged and mature mouse bladder sections. Aged sections had a higher number of *Jchain+* spot clusters than mature sections (Aged = 462, Young = 3).

